# The PMADS Project: A Longitudinal Multimodal Cohort Study to Understand Risk for Perinatal Mood and Anxiety Disorders

**DOI:** 10.64898/2026.04.10.717834

**Authors:** Noemí Rubau-Apa, Caroline Hayes, Ashley Francisco, Sage Rush, Hudah Rana, Mahima Islam, Lily Hunter, Laura Pritschet, Taylor Salo, Suneeta Senapati, Liisa Hantsoo, Deepak Indrakanti, Emily M. Beydler, Erica B. Baller, Ran Barzilay, Monica E. Calkins, Matthew Cieslak, John A. Detre, Sammy Dhaliwal, Mark A. Elliott, Hao Huang, Arielle S. Keller, C. Brock Kirwan, Rachel Kishton, Tyler M. Moore, Sara L. Kornfield, J. Cobb Scott, Manuel Taso, M. Dylan Tisdall, Arastoo Vossough, Lauren K. White, Kelly Zafman, Daniel H. Wolf, David R. Roalf, Sheila Shanmugan

## Abstract

**Background:** Perinatal mood and anxiety disorders (PMADs) are among the most common and consequential complications of pregnancy. The perinatal period is also characterized by profound hormonal fluctuations and large-scale brain plasticity. However, the mechanisms linking these neurobiological changes to psychiatric risk are poorly understood. Prospective, clinically informed studies are needed to identify quantitative biomarkers and clarify pathways linking perinatal neurobiology to PMADs risk.

**Methods:** This report describes the design of a prospective, longitudinal cohort study integrating multimodal neuroimaging, biofluid sampling, and deep clinical phenotyping to enable precision characterization of neurobiological trajectories of PMADs risk. Twenty-five individuals at elevated risk for PMADs will be recruited prior to conception and followed across six in-person timepoints spanning the menstrual cycle, pregnancy, and early postpartum, with additional remote follow-ups through the first postpartum year. Data collection includes high-resolution structural MRI, functional brain mapping using multi-echo resting-state fMRI, diffusion MRI, arterial spin labeling, ultra-high field MR-based techniques for measuring glutamate (GluCEST and ^1^HMRS), biofluid sampling, and comprehensive clinical, behavioral, and cognitive assessments. Structured clinical interviews assess categorical diagnoses while dimensional symptom measures capture heterogeneity and transdiagnostic features of perinatal psychopathology. Longitudinal analyses will model nonlinear trajectories of brain and symptom change across the perinatal period as well as evaluate whether preconception network features and menstrual cycle-related brain changes are associated with subsequent perinatal symptom emergence.

**Discussion:** This cohort study establishes a longitudinal, multimodal framework for investigating neurobiological changes across the transition to pregnancy in individuals at elevated risk for PMADs. By anchoring pregnancy-related brain changes to preconception and menstrual cycle-related variability within the same individuals, this study is designed to evaluate associations between preconception hormone sensitivity, pregnancy-induced neuroplasticity, and PMADs risk. The resulting dataset will provide a deeply phenotyped longitudinal resource for investigating brain-behavior relationships across the perinatal period. Findings are expected to inform future larger-scale studies aimed at advancing mechanistic understanding of PMADs, improving individualized risk stratification, and supporting development of personalized preventive and neuromodulatory interventions.

## Introduction

### Background

Mental health conditions are a current leading cause of maternal mortality in the United States, accounting for nearly 22% of pregnancy-related deaths (1,2). Perinatal mood and anxiety disorders (PMADs) are among the most common complications of pregnancy and childbirth, with perinatal depression alone affecting up to one in five birthing individuals (3). PMADs encompass depressive, anxiety, and related disorders with onset during pregnancy or within 12 months postpartum (3,4). Although prevalence has increased over the past decade, PMADs remain substantially under-recognized and undertreated. Their consequences are far-reaching, contributing to impaired maternal functioning, disrupted bonding and breastfeeding, elevated risk for chronic psychiatric illness, increased risk of obstetric complications such as preeclampsia and preterm birth, and adverse infant outcomes across socioemotional development, stress physiology, and sleep regulation (3). Early identification and effective intervention for PMADs are critical. Risk assessment for PMADs ideally occurs prior to conception in an effort to minimize overall exposures of both medications and untreated psychiatric symptoms (5). During preconception counseling, clinicians currently attempt to determine a patient’s risk for symptom recurrence during the perinatal period based on history and individual circumstances. However, quantitative neurobiological markers that could support precise risk assessment and guide clinical decisions, such as whether or when to initiate medications with unfavorable reproductive safety profiles, are lacking. Identifying such markers holds the potential to shift this paradigm, enabling more precise, personalized, and evidence-based guidance for patients. Understanding the neurobiological changes that unfold across the perinatal period, from preconception through the postpartum, is a critical step toward identifying such markers.

The perinatal period is characterized by profound hormonal fluctuations and large-scale neuroplasticity. Structural MRI studies consistently demonstrate gray matter volume and cortical thickness reductions across pregnancy, particularly within association networks such as the default mode, salience, and frontoparietal control systems, with partial recovery in the postpartum period (6–10). Resting-state fMRI studies reveal parallel reorganization of functional connectivity (11), pointing to plasticity in circuits supporting emotion regulation, threat appraisal, and caregiving (10). Recent work has begun to associate peripartum changes in limbic regions with depression symptoms and birth experience (12). Despite growing evidence that the perinatal period is marked by pronounced structural and functional brain reorganization, the field lacks a mechanistic understanding of how these changes relate to the onset, persistence, or remission of perinatal mood and anxiety symptoms. Notably, the regions that undergo the greatest neuroplastic changes across pregnancy overlap with neural circuitry implicated in mood and anxiety disorders (13), suggesting a potential pathway through which deviations from normative adaptation that reflect heightened neuroplasticity or increased vulnerability to adverse environmental influences may confer psychiatric risk in the context of preexisting risk factors or concurrent psychosocial stressors.

These gaps raise key questions: do individual differences in sensitivity to hormonal fluctuations shape how the brain adapts during the perinatal period, and is this heightened sensitivity observable prior to conception? One promising framework for addressing this question comes from considering pregnancy-related brain changes in the context of earlier, normative hormonal fluctuations across the menstrual cycle. Converging evidence across structural, functional, and molecular imaging demonstrates inter-individual differences in brain organization that are linked to hormone influences on microstructural and macrostructural processes (14,15). These findings suggest that gestational hormonal surges during pregnancy and abrupt withdrawal postpartum may differentially reorganize brain structure, cerebral perfusion, and functional architecture across individuals, acting as a risk factor for PMADs in individuals with latent neurobiological vulnerability (16,17). Importantly, such variability may already be detectable during normative hormonal fluctuations across the menstrual cycle, consistent with clinical observations that individuals with premenstrual dysphoric disorder (PMDD), premenstrual mood exacerbation, or sensitivity to exogenous hormones may be at elevated risk for perinatal depression (18–21) (Figure 1). While this literature suggests that sensitivity to normative hormonal fluctuations may index broader neurobiological vulnerability to reproductive mood disorders, no studies have used a within-person approach to link preconception hormonal sensitivity, perinatal brain remodeling, and PMADs symptoms.

**Figure 1.**
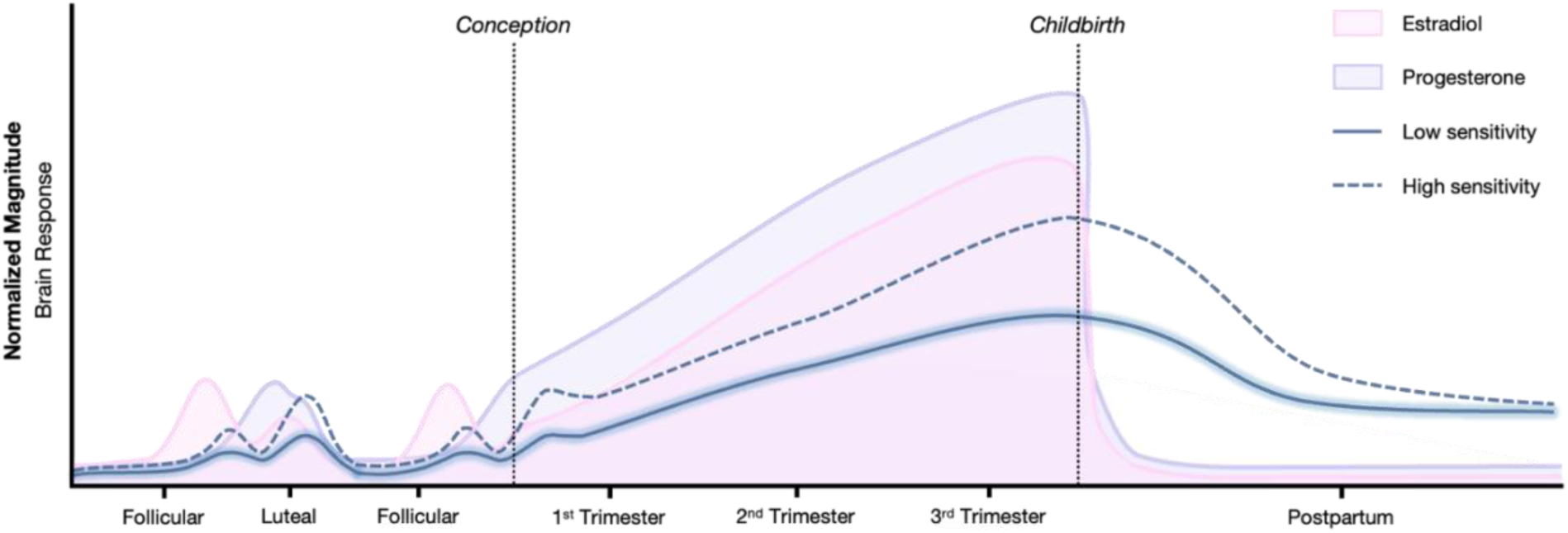
Conceptual Model of Hormonal Sensitivity–Driven Network Reorganization and Perinatal Vulnerability Estradiol (pink) and progesterone (purple) illustrate typical ovarian hormone trajectories across the menstrual cycle, pregnancy, and postpartum period. Solid and dashed blue lines represent hypothesized examples of individuals with relatively low versus high neural sensitivity to reproductive hormone fluctuations. Although hormonal concentrations follow broadly normative patterns, individuals may differ in the extent of neural or affective responses to these fluctuations. Such hormone sensitivity phenotypes may be observable during menstrual cycle transitions and may confer vulnerability to mood disturbance during greater endocrine shifts, such as during pregnancy or postpartum.

Recent advances in precision neuroimaging suggest that individualized mapping of functional neuroanatomy may better capture such inter-individual variability in hormone sensitivity that could be relevant to individual risk but that is obscured by group-level atlases (5). Precision functional mapping enables high-resolution characterization of personalized functional networks (PFNs) at the individual level, allowing for more reliable estimation of within-person patterns of connectivity and topography (22). Recent evidence demonstrates that PFNs provide stable and reproducible representations of brain organization that capture meaningful variation in cognition and psychopathology (23–26) and may serve as indicators of future risk for major depressive disorder (25). In the context of reproductive neuroendocrine transitions, this approach is particularly well-suited to capturing how individual differences in hormone sensitivity may manifest as distinct, person-specific trajectories of brain reorganization. Leveraging both longitudinal sampling as well as advanced acquisition strategies such as multi-echo fMRI (27,28), the present study aims to resolve individualized trajectories of network change across the perinatal period and to link these patterns to variation in hormonal sensitivity and PMAD risk.

In this study, we leverage high-quality structural, diffusion, perfusion, functional, and molecular data to capture complementary aspects of brain organization and plasticity across the perinatal transition. This multimodal, deeply phenotyped, and densely sampled approach reflects a broader shift toward precision neuroscience, enabling more accurate characterization of individual trajectories of brain change (29), an advancing over prior designs with limited temporal resolution across gestation. We aim to identify preconception markers of PMADs symptoms by evaluating whether individual differences in person-specific imaging measures across the menstrual cycle are associated with later vulnerability to perinatal depression and anxiety. We hypothesize that greater menstrual cycle-related differences in imaging metrics between the follicular and luteal phases will be associated with increased risk for developing depressive and anxiety symptoms during the perinatal period. We also aim to identify brain changes associated with the emergence of depressive and anxiety symptoms during pregnancy and the postpartum period. We hypothesize that the emergence of PMADs symptoms will be associated with deviations from normative perinatal trajectories of structural and functional remodeling, particularly in association cortex.

## Methods

### Study Design

This is a prospective, longitudinal, observational study. Data collection includes a combination of behavioral, cognitive and clinical assessments, biofluid collection, and 3T and 7T magnetic resonance imaging (MRI). Neuroimaging measures include structural MRI, multi-echo functional MRI (fMRI), diffusion MRI, arterial spin-labeled perfusion (ASL) MRI, glutamate-weighted chemical exchange saturation transfer (GluCEST), and proton magnetic resonance spectroscopy (^1^HMRS). As shown in Figure 2, the study follows a structured sequence of assessments across key phases.

**Figure 2.**
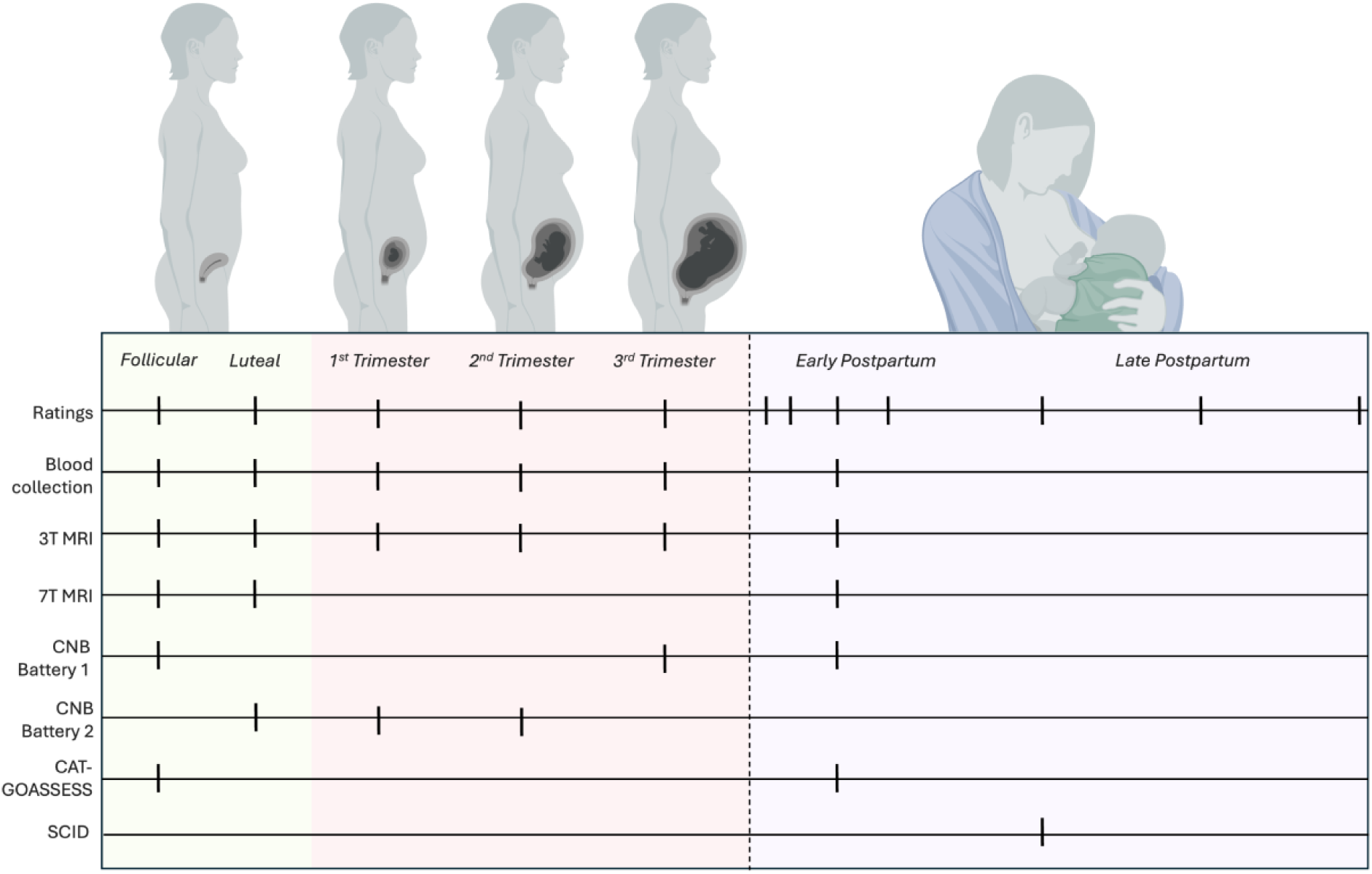
Study Design and Assessment Timeline Prospective longitudinal study design spanning the preconception period (yellow), pregnancy (red), and the first postpartum year (purple). Participants complete structured assessments during the follicular and luteal phases prior to conception, three trimester-based visits during pregnancy, and serial follow-up assessments postpartum. Tick marks indicate timepoints at which behavioral, clinical, biological, cognitive, and neuroimaging measures are collected. 3T and 7T MRI scans may be completed during a single visit or split across two visits. If MRI sessions occur more than 24 hours apart from biofluid and ratings, repeat collection occurs. Shaded regions denote reproductive stage. The dashed vertical line marks delivery. Abbreviations: CNB, Penn Computerized Neurocognitive Battery; CAT-GOASSESS, Computerized Adaptive Test version of the GOASSESS; SCID, Structured Clinical Interview for DSM-5. *Created in BioRender. Rubau Apa, N.* (*2026*) https://BioRender.com/9bnr0sf

The study team first identifies and recruits participants using a secure RedCap screening form. Participants undergo a brief phone screen followed by a more in-depth virtual screening visit. Eligible participants who enroll in the study are then assessed preconception, during pregnancy, and postpartum. Participants complete preconception study visits in both the follicular and luteal phases of their menstrual cycle. Follicular study visits occur between cycle days 3-7 to capture a low-hormone baseline state characterized by relatively low levels of estradiol and progesterone. Luteal study visits occur 8-12 days following ovulation, with ovulation timing determined using at-home urine LH surge testing to ensure precise capture of the mid-luteal, high progesterone state. This staging enables the examination of neurobiological variation in relation to endogenous hormone fluctuations prior to pregnancy. Participants are assessed three times across pregnancy to allow for prospective tracking of mood, brain function, and biological measures as pregnancy progresses. Assessments during pregnancy correspond to roughly one study visit in each trimester, with flexibility for the first visit to occur in the late first or early second trimester to mitigate drop out due to normal pregnancy-related nausea and vomiting as well as early pregnancy loss. Participants who do not conceive within 9 months of completing preconception assessments will be invited to repeat the follicular study visit, thereby serving as a longitudinal non-pregnant control arm. In the postpartum period, participants complete one to two in-person study visits within the first 5 months after delivery, one virtual visit at approximately 7 months postpartum, and seven asynchronous follow-ups over the next year, enabling assessment of psychological and biological trajectories across the first postpartum year.

### Participants

This study aims to recruit 25 female individuals between the ages of 25 and 40 who are at risk for developing perinatal mood and anxiety disorders (PMADs). Participants entering the study during the preconception period must be nulliparous and either actively trying to conceive or planning to start within the next three months. Menstrual cycle regularity is required for participation in luteal phase-specific components and is assessed via self-report questionnaires during screening. Regular cycles are defined as meeting all of the following criteria in the absence of current systemic hormonal contraceptive use: average cycle length between 24–38 days, cycle length variability ≤7 days over the last 12 cycles, and bleeding duration ≤8 days. Participants must endorse at least one of the following risk factors for PMADs to be eligible for inclusion: (1) current premenstrual dysphoric disorder (PMDD), premenstrual exacerbation of underlying mood disorder (PME), or other significant menstrual-related mood symptoms or (2) current or past mood sensitivity to hormonal contraception or fertility treatments, including individuals who discontinued use due to adverse mood effects. Risk factor status endorsed on a prescreen survey is assessed via second-level screening with the Premenstrual Symptoms Screening Tool (PSST) (13), medical record review and the Reproductive Mental Health History Questionnaire - G0 (Gravida 0) Version, a study-specific instrument designed for nulliparous women developed to assess preconception history of mood sensitivity to reproductive hormonal fluctuations (see Supplementary Material 1). This questionnaire assesses prior diagnoses of mood and anxiety disorders; patterns of menstrual-related mood symptoms and associated functional impairment; family history of perinatal depression; and mood responses to exogenous hormonal exposures, including hormonal contraception and fertility treatments. This detailed assessment enables characterization of exposure-specific symptom patterns that may index vulnerability to PMADs. We anticipate that this questionnaire may evolve as we test and validate it during the course of the study.

Exclusion criteria include major neurological conditions, chronic opiate use, chronic pain, active post-traumatic stress disorder (PTSD), borderline personality disorder (BPD), primary psychotic disorders, and active substance use, including frequent alcohol use (≥8 drinks per week or alcohol consumption on ≥5 days per week), weekly tetrahydrocannabinol (THC) use, or any nicotine use. Participants are instructed to abstain from alcohol for 48 hours prior to each MRI visit to reduce acute confounding effects of alcohol on neuroactive steroids. Participants are also asked to abstain from high dose biotin supplements for 72 hours prior to blood draws to prevent interference with sex steroid hormone assays (30). Participants with a history of pregnancy beyond 12 weeks are excluded to minimize the potential confounding effects of prior pregnancy-related neuroplastic changes (31). Diminished ovarian reserve (unless under care of reproductive endocrinology and infertility providers), early menopause, unilateral oophorectomy, or hysterectomy are also exclusionary. Body mass index (BMI) outside the range of 17-27 may be exclusionary unless menstrual regularity or care by a reproductive endocrinologist is documented. Additional exclusion criteria include any contraindications to MRI, medical complexity, and inability or unwillingness to attend in-person study visits.

Primary recruitment methods include MyChart messaging, a feature of Penn Medicine’s electronic medical record system (EPIC); patients who meet basic demographic eligibility criteria (e.g., age, sex, and obstetric history) and who are actively seeking care related to preconception counseling, assisted reproductive technologies, or removal of a long-acting reversible contraceptive method are identified. Additional recruitment strategies include recontacting individuals who previously participated in research within the Department of Psychiatry and distributing flyers in key clinical and public locations across the University of Pennsylvania and the broader Philadelphia community. Physician referrals further support recruitment. All study procedures have been approved by the Institutional Review Board of the University of Pennsylvania, and all participants provide written informed consent prior to participation.

### Behavioral and Clinical Measures

#### Self-Report Scales

Self-report data are collected across multiple symptom domains to capture both primary and comorbid features of perinatal mental health. Although perinatal depression is the primary symptom domain of interest given its high prevalence in this population (3), inclusion of additional symptom domains allow us to assess comorbidity and explore prognostic secondary analyses of other disorders. This multidomain approach is intended to capture the heterogeneity and complexity of perinatal psychopathology. Table 1 summarizes the questionnaire battery, including the domains assessed and their administration timepoints.

**Table 1.**
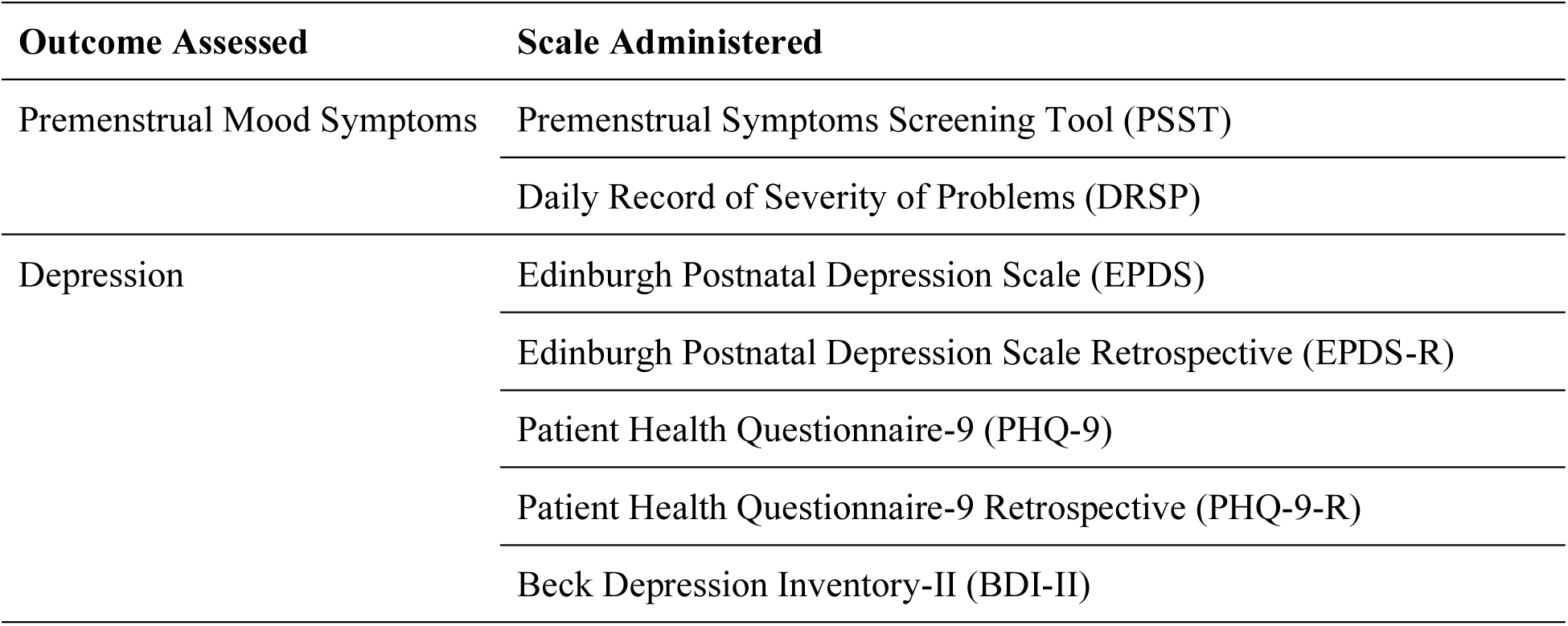
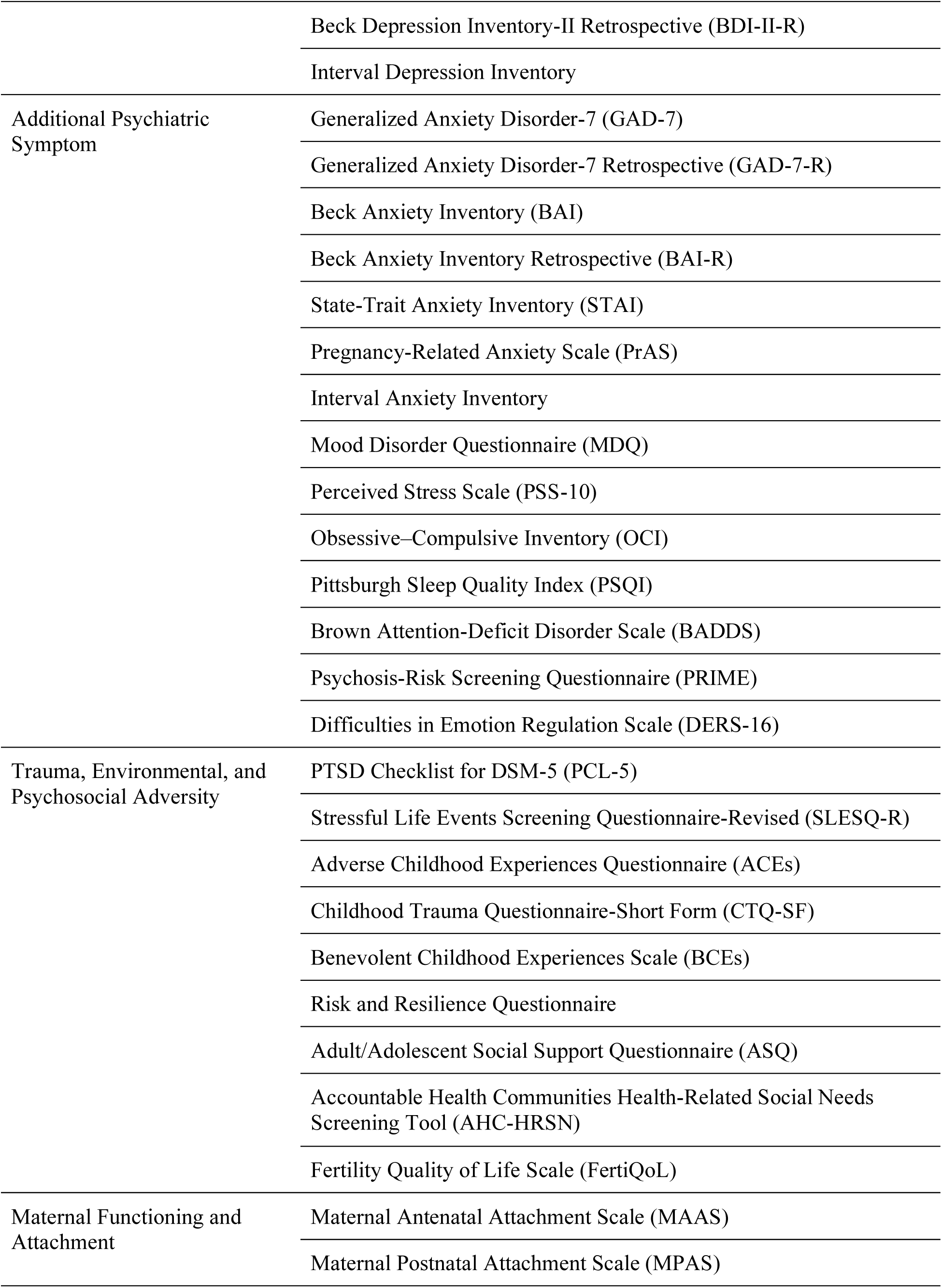
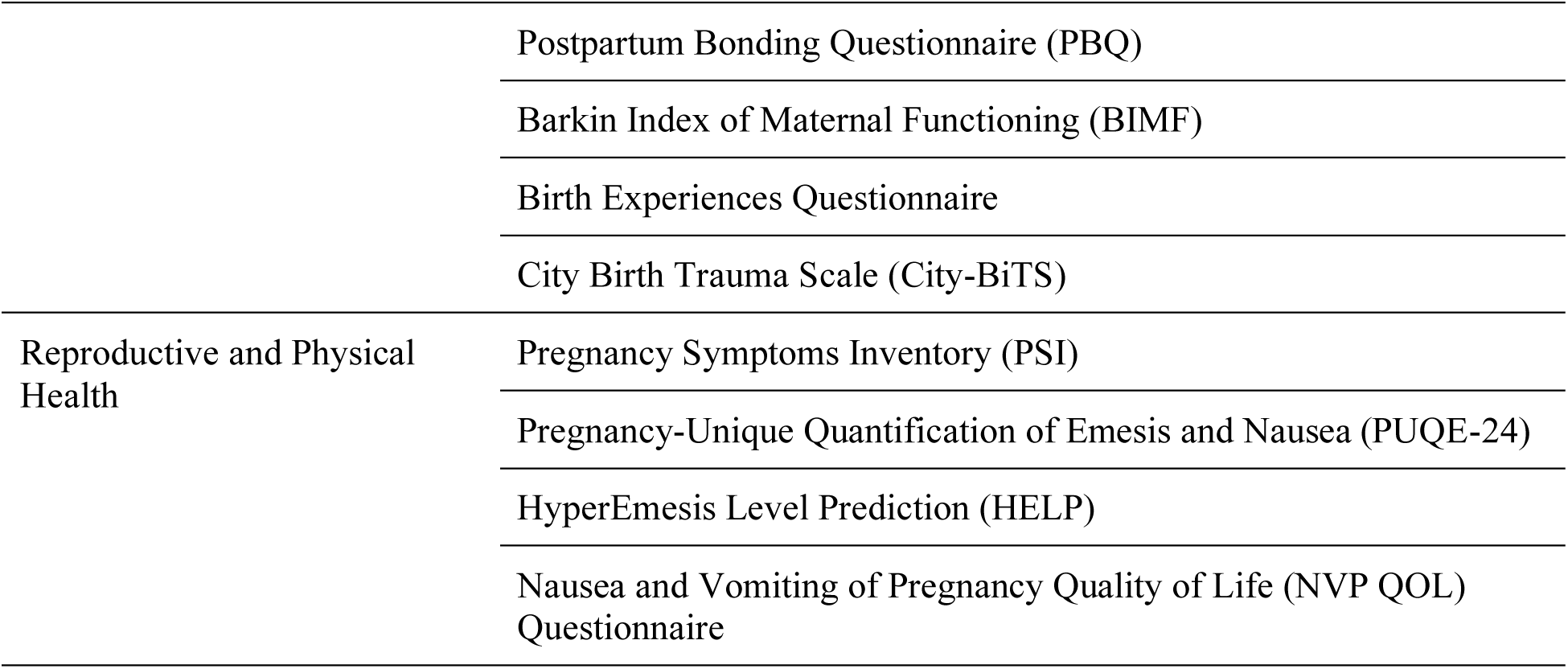
Scales and outcomes assessed.

##### Premenstrual Mood Symptoms

Menstrual-cycle related mood symptoms are assessed using the Premenstrual Symptoms Screening Tool (PSST) (32). Although prospective daily symptom tracking across at least two menstrual cycles is considered the gold standard for diagnosing PMDD, retrospective instruments such as the PSST have demonstrated good sensitivity and clinical utility in identifying individuals with clinically significant premenstrual mood symptoms (33). To prospectively assess menstrual-cycle related mood symptoms, participants complete the Daily Record of Severity of Problems (DRSP) daily for two full menstrual cycles using the myCap app. The DRSP is a validated, prospective symptom-tracking measure that provides multi-faceted characterization of mood, behavioral, physical, and functional symptoms, enabling assessment of day-to-day depressive symptom variability and cycle-related fluctuation (34). Participants are sent a text message reminder at approximately 7 PM each day prompting them to complete the DRSP prior to midnight. Completion is monitored so that study staff can reach out to participants who miss two consecutive days of DRSP entries to discuss alternative reminder strategies (e.g., changing the time of the reminder) or completion methods (e.g., phone calls) to promote daily adherence.

##### Depression Measures

Depressive symptoms are assessed using multiple complementary scales across preconception, pregnancy and postpartum. The Edinburgh Postnatal Depression Scale (EPDS) (35) is a 10-item questionnaire designed to assess perinatal depressive symptoms over the past seven days. Items are scored on a 4-point Likert scale (0–3), yielding a total score range of 0–30, with higher scores indicating more severe depressive symptoms. The Patient Health Questionnaire-9 (PHQ-9) (36) is administered as a general measure of depression severity. This 9-item instrument assesses the frequency of depressive symptoms over the past two weeks on a 0-3 scale, with a total score range of 0-27. To capture a broader spectrum of depressive symptomatology, the Beck Depression Inventory-II (BDI-II) (37) is also included to assess cognitive, affective, and somatic features of depression, scored from 0-3 per item to yield a range of 0–63. In addition to prospective assessments, the Interval Depression Inventory (Supplementary Material 2) as well as retrospective versions of the EPDS, PHQ-9, and BDI-II are administered to reconstruct symptom trajectories between study visits and enhance detection of mood symptoms and proactive treatment occurring outside scheduled assessment windows. Retrospective versions of the EPDS, PHQ-9, and BDI-II retain identical item content to the validated instruments but instruct participants to rate symptoms relative to the interval since their prior study visit. Similarly, retrospective versions of the EPDS, PHQ-9, and BDI-II are used to assess depressive symptoms associated with significant decreases in milk supply or weaning.

##### Additional Psychiatric Symptom Measures

A multidomain set of instruments is included to capture common psychiatric comorbidities in the perinatal period. Anxiety symptoms are assessed with the Generalized Anxiety Disorder-7 (GAD-7) (38), a 7-item self-report measure evaluating generalized anxiety over the past two weeks; the Beck Anxiety Inventory (BAI) (39), a 21-item scale measuring somatic and cognitive aspects of anxiety; the State-Trait Anxiety Inventory (STAI) (40), which distinguishes between state-dependent and trait-based anxiety; and the Pregnancy-Related Anxiety Scale (PrAS) (41), a pregnancy-specific instrument targeting concerns about maternal and fetal health. To supplement these instruments, the Interval Anxiety Inventory (Supplementary Material 3) as well as a retrospective version of the GAD-7 and the BAI are administered during pregnancy and postpartum to capture anxiety symptoms that emerged, remitted, or were treated between study visits. Similarly, a retrospective version of the GAD-7 is used to assess anxiety symptoms associated with periods of significant decreases in milk supply or weaning.

Broader psychiatric and behavioral domains are also measured. The Mood Disorder Questionnaire (MDQ) (42) screens for bipolar spectrum symptoms, while the Perceived Stress Scale (PSS-10) (43) captures perceived stress over the past month. The Obsessive–Compulsive Inventory (OCI) (44) is a self-report instrument for obsessive-compulsive symptoms, and the Pittsburgh Sleep Quality Index (PSQI) (45) provides a global measure of sleep quality and disturbances. The Brown Attention-Deficit Disorder Scale (BADDS) for adults assesses subjective executive function difficulties, and the Psychosis-Risk Screening Questionnaire (PRIME) (46) screens for subthreshold psychotic symptoms.

Emotion regulation difficulties are assessed using the 16-item Difficulties in Emotion Regulation Scale (DERS-16) (47), a validated brief measure of multidimensional regulation deficits. Emotion regulation has been robustly associated with perinatal depressive symptoms in large cohort studies (48), supporting its inclusion as a transdiagnostic vulnerability process in PMADs.

##### Trauma, Environmental, and Psychosocial Measures

Several instruments capture trauma exposure, psychosocial adversity, and protective factors relevant to perinatal mental health. Trauma symptoms and histories are assessed with the PTSD Checklist for DSM-5 (PCL-5), a 20-item measure of post-traumatic stress symptoms administered after birth, and the Stressful Life Events Screening Questionnaire-Revised (SLESQ-R) (49), which provides a checklist of exposure to potentially traumatic events. The Adverse Childhood Experiences Questionnaire (ACEs) (50) and Childhood Trauma Questionnaire-Short Form (CTQ-SF) (51) capture early-life adversity, while the Benevolent Childhood Experiences Scale (BCEs) provides a complementary assessment of protective childhood factors.

Environmental and psychosocial factors are measured with the Risk and Resilience Questionnaire (52), which captures coping resources and vulnerability markers, the Adult/Adolescent Social Support Questionnaire (ASQ) (53), and the Accountable Health Communities Health-Related Social Needs Screening Tool (AHC-HRSN) (54), which screens for social determinants of health such as housing, food, and interpersonal safety. The Fertility Quality of Life Scale (FertiQoL) (55) is administered to assess the impact of fertility and reproductive health challenges on psychosocial well-being. Although substance use is exclusionary at baseline, participants complete the Timeline Follow-Back (TLFB) to assess use of alcohol or other substances that may impact measures of interest (e.g. neuroactive steroids) in the relevant time interval prior to MRI scans.

##### Maternal Functioning and Attachment

Measures of maternal role adaptation and mother-infant relationships are collected across pregnancy and postpartum. The Maternal Antenatal Attachment Scale (MAAS) (56) is a self-report instrument assessing the strength and quality of emotional attachment to the fetus during pregnancy, while the Maternal Postnatal Attachment Scale (MPAS) (57) parallels this assessment in the postpartum period, evaluating maternal affect, pleasure in interaction, and absence of hostility toward the infant. Bonding after delivery is further assessed with the Postpartum Bonding Questionnaire (PBQ) (58), a 25-item measure that identifies difficulties in establishing a positive mother-infant relationship.

To evaluate functional outcomes, the Barkin Index of Maternal Functioning (BIMF) (59) is administered, a 20-item scale designed to measure maternal competence and well-being in the postpartum period. Complementary domains are captured with the Birth Experiences Questionnaire (60), which assesses subjective appraisals of the delivery process, and the City Birth Trauma Scale (City-BiTS) (61), which assesses trauma specifically related to childbirth.

##### Reproductive and Physical Health Measures

A set of instruments capture gynecologic, reproductive, and somatic health domains relevant to perinatal mental health. The Pregnancy Symptoms Inventory (PSI) (62) assesses physical symptoms experienced during pregnancy. Nausea and vomiting of pregnancy (NVP) are assessed using the Pregnancy-Unique Quantification of Emesis and Nausea (PUQE-24) (63) scale and the HyperEmesis Level Prediction (HELP) score (64), which quantify symptom severity and risk for clinically significant hyperemesis. In addition, the Nausea and Vomiting of Pregnancy Quality of Life (NVP-QOL) questionnaire (65) is administered to capture the functional impact of symptoms on daily life.

In addition to standardized scales, participants complete study-specific forms to provide detailed medical and reproductive histories, medication use, and supplements. Data from these forms are supplemented by electronic medical record (EMR) review, allowing integration of self-reported and clinically verified health information.

#### Clinical Assessment

Participants undergo a structured clinical interview at a 7 months postpartum visit to determine current and lifetime diagnoses of psychiatric disorders based on DSM-5 criteria. The Structured Clinical Interview for DSM-5 (SCID-5) (66) is administered to assess major diagnostic domains relevant to internalizing and related psychopathology. Modules administered include mood, anxiety, obsessive-compulsive and related, trauma- and stressor-related, psychotic, substance use disorders, as well as insomnia disorder. To minimize participant burden, modules assessing externalizing, feeding and eating disorders, and other diagnostic categories are omitted. This timing is selected to capture the majority of perinatal symptom onset, which most commonly occurs during pregnancy or within the first 6 months postpartum (67).

In addition to this structured interview that yields categorical DSM diagnoses, participants also complete the Computerized Adaptive Testing version of the GOASSESS (CAT-GOASSESS) (68), a clinical assessment that provides a dimensional characterization of lifetime psychopathology. Adapted from the structured GOASSESS screening interview (69), the CAT-GOASSESS assesses five transdiagnostic domains: internalizing symptoms (e.g., mood and anxiety), phobias, externalizing symptoms, psychosis, and personality disorder symptoms. By assessing symptoms dimensionally rather than categorically, the CAT-GOASSESS is well suited to capture subthreshold symptomatology and high rates of psychiatric comorbidity, which are common in population-based samples and may be obscured by categorical, diagnosis-focused assessments. The CAT-GOASSESS is optimized for brief administration, typically requiring approximately 8 minutes and minimal proctoring (68). Item-level data from the CAT-GOASSESS will be used to calculate continuous factor scores for transdiagnostic domains, as well as an overall score reflecting general psychopathology (68). Table 2 summarizes the domains and associated psychopathology captured by the CAT-GOASSESS.

**Table 2.** CAT-GOASSESS.

| Domain | Example associated psychopathology |
| --- | --- |
| Mood and anxiety | Major depressive episode, mania/hypomania |
| Phobias | Agoraphobia, specific phobia |
| Externalizing disorders | Attention deficit/hyperactivity, oppositional defiant, conduct |
| Psychosis | Psychosis, prodromal psychosis |
| Personality | Negative affect, detachment, antagonism, disinhibition, psychoticism |

#### Computerized Neurocognitive Battery

Cognitive functioning will be assessed using a customized version of the Penn Computerized Neurocognitive Battery (CNB) (70) given that cognitive dysfunction is a core feature of depression (71) and has been shown to change across the perinatal period (72), although the specific domains affected, their longitudinal trajectories, and their neurobiological correlates remain poorly understood. Each administration of the CNB takes approximately one hour and is completed under supervision by trained research staff. Tasks are drawn from validated instruments that assess neurobehavioral domains including executive functioning, episodic memory, complex reasoning, social cognition, and sensorimotor processing (73). Two CNB batteries are administered in this study.

Battery 1 includes the Psychomotor Vigilance Test (PVT), Digit Symbol Test, Short Penn Continuous Performance Test (sPCPT), Short Letter-N-Back (SLNB2), and the Penn Abstraction, Inhibition, and Working Memory Task (AIM). We also use Adaptive CNB tasks (74). The adaptive battery includes the Penn Emotion Recognition Test (ER40), Age Differentiation Test (ADT), Measured Emotion Differentiation Test (MEDF), Penn Matrix Reasoning Test (PMAT), Line Orientation Test (PLOT), Short Verbal Reasoning Test (SPVRT) (Table 3a). Administration is guided by standardized protocols and supported by an internal training system. Participants complete Battery 1 at three study visits: follicular, third trimester and first postpartum visit. Additionally, participants complete Battery 2 including the PVT and Digit Symbol Test tasks (Table 3b), which have limited practice effects, at each study visit.

**Table 3.** CNB.

**a. Battery 1**
| Test Name | Domain Assessed |
| --- | --- |
| Psychomotor Vigilance Test | Vigilant Attention |
| Digit Symbol Test | Processing Speed & Working Memory |
| Short Penn Continuous Performance Test | Sustained Attention |
| Short Letter-N-Back; 2-, and 3-Back Versions | Working Memory |
| Penn Abstraction, Inhibition and Working Memory Task | Abstraction/Inhibition |
| Penn Emotion Recognition Test | Emotion Identification |
| Age Differentiation Test | Age Differentiation |
| Measured Emotion Differentiation Test | Emotion Intensity Differentiation |
| Penn Matrix Reasoning Test | Non Verbal Reasoning |
| Penn Line Orientation Test | Spatial Processing |
| Short Penn Verbal Reasoning Test | Language Reasoning |

**b. Battery 2**
| Test Name | Domain Assessed |
| --- | --- |
| Psychomotor Vigilance Test | Vigilant Attention |
| Digit Symbol Test | Processing Speed & Working Memory |

### Hormonal and Biological Sampling

#### Blood collection

Venous blood is collected at all in-person study visits spanning preconception, pregnancy, and postpartum. Blood draws include silicone-coated serum tubes with clot activator for collection of serum, as well as ethylenediamine tetraacetic acid (EDTA) lavender-top tubes for collection of plasma and whole blood. All blood samples are collected via standard phlebotomy and processed according to standardized procedures. Samples that are not immediately assayed are stored at –80 °C until analysis.

EDTA tubes are processed as soon as possible, no more than 30 minutes after collection. Serum is processed at 4 °C within 30 minutes of collection. Serum samples not processed within 40 minutes are placed in a cooler with ice, wrapped in a paper bag to avoid erythrocyte lysis due to direct contact of tubes with ice. After centrifuging, samples are kept at 2–8°C (on ice) while handling and aliquoting into cryovials. Cryovials are placed in a cooler with dry ice and transferred to a -80°C freezer for storage.

Serum samples will be used to quantify allopregnanolone and related neuroactive steroid metabolites as well as for quantification of sex steroid and gonadotropic hormones, including estradiol, progesterone, luteinizing hormone, and follicle-stimulating hormone. Plasma aliquoted from EDTA tubes and/or serum will be used for high-dimensional proteomic profiling using SomaScan or a comparable affinity-based proteomic platform. Whole blood collected in EDTA tubes will be stored at -80°C for future genomic and epigenetic analyses (e.g., genetic variation, DNA methylation). A complete blood count (CBC) will be obtained in a clinical laboratory from whole blood at each 3T MRI study visit to account for pregnancy-related changes in hematocrit in analyses of ASL data.

All specimens are linked to clinical metadata (visit, reproductive stage, medication status) and assayed in batches/one batch to minimize inter-assay variability.

### Neuroimaging acquisition protocols

#### Participant positioning

During MRI, participants are positioned with a 30° left lateral tilt (LLT) to minimize compression of the inferior vena cava (IVC) during pregnancy. In both clinical and research settings, brain MRI acquisition is typically performed in the supine position; however, prolonged supine positioning during later pregnancy can result in compression of the IVC by the gravid uterus (75,76). IVC compression reduces venous return to the heart and is associated with decreased cardiac output and stroke volume in the supine position relative to left lateral positioning (76–78), even in the absence of overt symptoms due to compensatory autonomic mechanisms (79,80). Although most pregnant individuals can tolerate brief periods in the supine position, it remains unclear whether subclinical alterations in cardiac output and venous return may influence neuroimaging metrics sensitive to blood oxygenation or cerebral perfusion (81,82).

In addition to potential physiological confounds, prolonged supine positioning may increase discomfort and anxiety in pregnant participants, as individuals are commonly counseled to avoid the supine position in later gestation due to concerns regarding aortocaval compression (75,83), despite more recent evidence suggesting that short-duration supine positioning in monitored settings does not independently increase stillbirth risk (75). This concern is especially relevant in the current study, which enrolls individuals at elevated risk for PMADs, many of whom have comorbid anxiety. To minimize potential confounding effects of positioning on imaging measures, reduce participant discomfort, and avoid introducing anxiety-related topics to participation, all participants are imaged in a standardized 30° LLT position. This position has been shown to significantly increase IVC volume and improve hemodynamic parameters compared to the supine position (76,84,85), while allowing head positioning compatible with standard preprocessing pipelines. Although a full 90° left lateral decubitus position would likely maximize participant comfort and relief of IVC compression, this position is not compatible with modern multi-channel head coils with tight enclosures and would introduce substantial deviation from conventional acquisition geometry.

In this study, positioning is initiated by placing participants briefly in a full left lateral decubitus position (approximately 90°) for 90 seconds to facilitate displacement of the gravid uterus off the midline and relieve aortocaval compression (76,86). Participants are then transitioned onto a 30° foam wedge positioned along the right side of the body to maintain a sustained LLT of the body. To minimize potential lateralizing effects of neck rotation, small cushions and pillowcases are used to ensure that the head and neck are stabilized in a position that is aligned with the body. The imaging field of view is not angled to compensate for head tilt due to impacts on preprocessing pipelines; instead, any deviation from standard axial orientation is addressed during preprocessing using validated pipelines.

#### 3T MRI Protocol

All 3T imaging is performed on a Siemens MAGNETOM 3T Prisma FIT running software version NXVA60 and using the vendor’s 32-channel head coil. The acquisition protocol includes structural MRI, multi-echo resting-state fMRI, diffusion MRI, and arterial spin-labeled perfusion MRI (Table 4). All sequences are performed in Normal Mode (whole-body SAR ≤ 2 W/kg), and a given sequence does not proceed if the system indicates potential peripheral nerve stimulation.

**Table 4.** 3T MRI Imaging parameters.

| Sequence | Slices | Resolution (mm) | TR (ms) | TE (ms) | TI (ms) | Flip Angle (°) | Multi Band Accel |
| --- | --- | --- | --- | --- | --- | --- | --- |
| anat-T1w_MEMPRAGE | 176 | $1.0 \times 1.0 \times 1.0$ | 2500 | 1.79, 3.71, 5.63, 7.55 | 1070 | 8 | — |
| fmap-epi_acq-func_dir-AP | 72 | $2.0 \times 2.0 \times 2.0$ | 14000 | 24.8, 69.4, 114.0, 158.6 | — | 90 | — |
| fmap-epi_acq-func_dir-PA | 72 | $2.0 \times 2.0 \times 2.0$ | 14000 | 24.8, 69.4, 114.0, 158.6 | — | 90 | — |
| func-bold_task-rest_dir-AP_run-01 | 72 | $2.0 \times 2.0 \times 2.0$ | 1761 | 16.8, 45.9, 75.0, 104.1 | — | 68 | 6 |
| func-bold_task-rest_dir-PA_run-02 | 72 | $2.0 \times 2.0 \times 2.0$ | 1761 | 16.8, 45.9, 75.0, 104.1 | — | 68 | 6 |
| dwi-dwi_acq-HBCD75_dir-AP | 87 | $1.7 \times 1.7 \times 1.7$ | 4800 | 93 | — | 78 | 3 |
| dwi-dwi_acq-HBCD75_dir-PA | 87 | $1.7 \times 1.7 \times 1.7$ | 4800 | 93 | — | 78 | 3 |
| perf-asl | 44 | $3.0 \times 3.0 \times 3.0$ | 5000 | 9.61 | — | 21.5, 75 | — |

High-resolution anatomical images are acquired using a T1-weighted MEMPRAGE sequence (87,88) with embedded volumetric navigators (nNav) (TR = 2500 ms; TE = 1.79, 3.71, 5.63, and 7.55 ms; TI = 1070 ms; flip angle = 8°; voxel size = 1.0 × 1.0 × 1.0 mm³; 176 sagittal slices; GRAPPA acceleration factor = 2; acquisition time = 6:09 min).

Resting-state functional data are collected with a multi-echo echo-planar imaging (EPI) (89) sequence in both anterior-posterior and posterior-anterior phase-encoding directions to enable distortion correction, with additional field map acquisitions collected to support susceptibility distortion correction (TR = 1761 ms; TEs = 16.8, 45.9, 75.0, and 104.1 ms; flip angle = 68°; voxel size = 2.0 × 2.0 × 2.0 mm³; 72 transversal slices; multiband factor = 6; 400 volumes per run; acquisition time = 12:30 min per run).

Diffusion images are acquired in both phase-encoding directions using a multi-band EPI sequence (TR = 4800 ms; TE = 93 ms; flip angle = 78°; voxel size = 1.7 × 1.7 × 1.7 mm³; 87 transversal slices; multiband factor = 3; 75 diffusion directions; b = 0 and 3000 s/mm²; acquisition time = 6:24 min per acquisition direction) (90).

Cerebral perfusion is measured with unbalanced pseudo-continuous arterial spin labeling (pCASL) using background suppression and a 3D stack-of-spirals readout (TR = 5000 ms; TE = 9.61 ms; labeling flip angle = 21.5°; refocusing flip angle = 75° voxel size = 3.0 × 3.0 × 3.0 mm³; labeling duration = 1800 ms; post-labeling delay = 1800 ms; 44 slices; 8 control/label pairs; acquisition time = 4:24 min).

#### 7T MRI Protocol

Enrollment in the 7T component of the protocol is optional, and participants may enroll in the remaining protocol without completing 7T imaging. Only participants who defer attempts to conceive for one menstrual cycle complete the 7T luteal MRI to ensure that 7T imaging is not conducted during pregnancy, given limited safety data at ultra-high field strengths (91). All ultra–high-field imaging is performed on a Siemens MAGNETOM 7T Terra system running software version VE12U-SP01 and using the vendor’s 32-channel head coil. The protocol integrates high-resolution structural MRI with multimodal glutamatergic-sensitive techniques, including chemical exchange saturation transfer (CEST) imaging and ¹HMRS (Table 5).

**Table 5.**
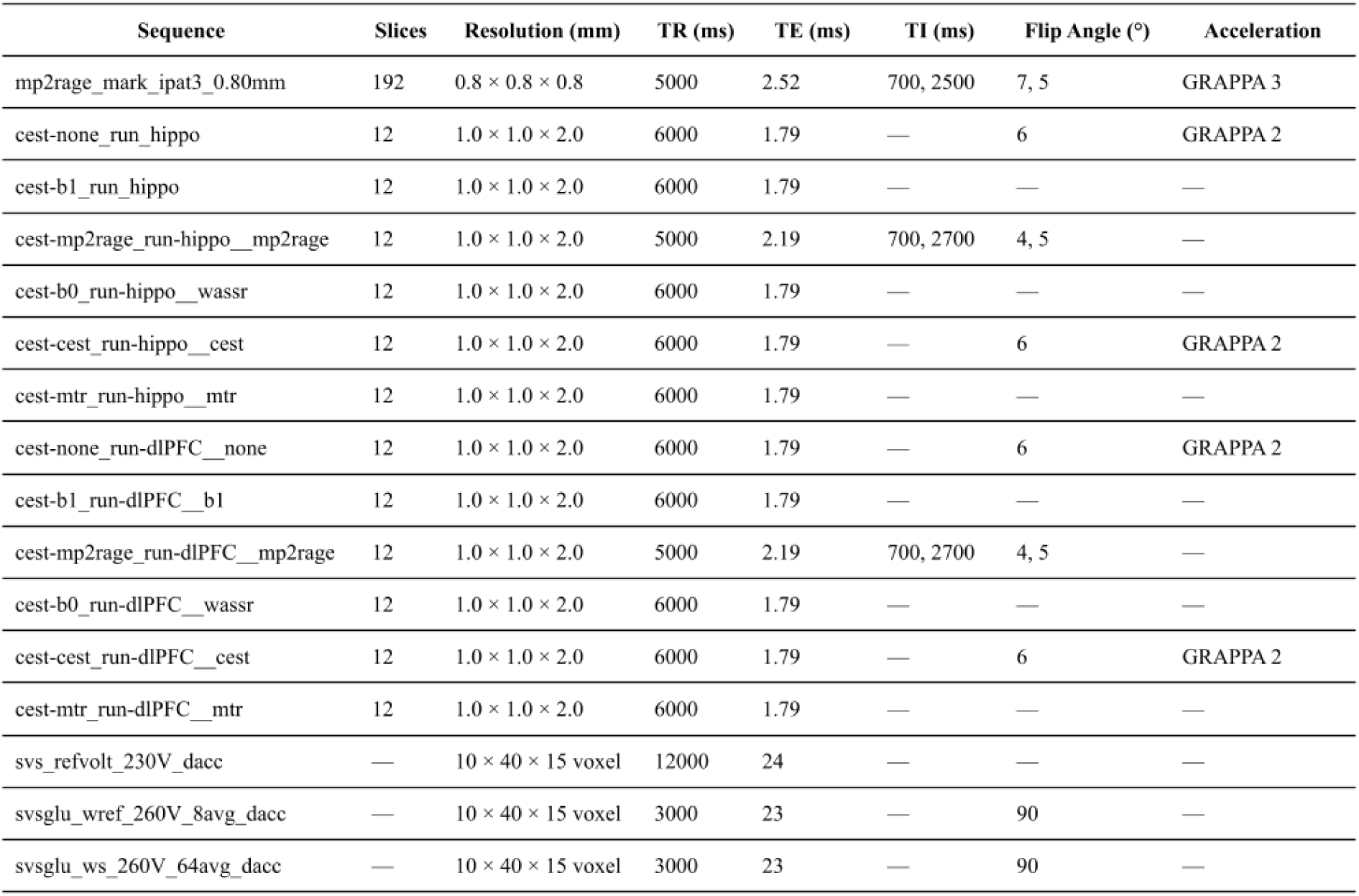
7T MRI Imaging parameters.

T1-weighted structural images are acquired at 7T using an MP2RAGE sequence to support high-resolution cortical and subcortical morphometric analyses as well as quantitative T1 mapping (92). A whole-brain MP2RAGE acquisition is collected with isotropic submillimeter resolution (0.8 × 0.8 × 0.8 mm³; TR = 5000 ms; TE = 2.52 ms; TI1 = 700 ms; TI2 = 2500 ms; flip angles = 7° and 5°; 192 slices; GRAPPA acceleration factor = 3; acquisition time = 7 min), optimized for segmentation and surface-based reconstruction.

GluCEST imaging (93) is performed in the hippocampus and dorsolateral prefrontal cortex (DLPFC) using region-targeted 3D slab-selective gradient-echo acquisitions. For both regions, the primary CEST acquisition is acquired with voxel size = 1.0 × 1.0 × 2.0 mm³, TR = 6000 ms, TE = 1.79 ms, flip angle = 6°, GRAPPA acceleration factor = 2, 12 slices per slab, FoV read = 240 mm, and phase resolution = 100%. The GluCEST acquisition samples frequency offsets from 1.8 to 4.2 ppm in 0.3-ppm steps. To correct for transmit and field inhomogeneities, separate B1 mapping and WASSR acquisitions are collected in the same field of view. The B1 map is acquired with the same geometric parameters as the CEST scan (TR = 6000 ms, TE = 1.79 ms, voxel size = 1.0 × 1.0 × 2.0 mm³) across 3 measurements. The WASSR acquisition is also matched in geometry (TR = 6000 ms, TE = 1.79 ms, voxel size = 1.0 × 1.0 × 2.0 mm³) and samples offsets from 0.0 to 1.5 ppm in 0.15-ppm steps, with saturation duration = 200 ms and preparation B1 = 20 Hz. Additional magnetization transfer reference scans are acquired from 20 to 100 ppm, and region-matched MP2RAGE images are collected for anatomical localization and T1 mapping within the CEST field of view (voxel size = 1.0 × 1.0 × 2.0 mm³, TR = 5000 ms, TE = 2.19 ms, TI1 = 700 ms, TI2 = 2700 ms, flip angles = 4° and 5°). All participants are registered to the same template for CEST acquisition at their first 7T study visit. At subsequent visits, Imscribe (cmroi.med.upenn.edu/imscribe) is used to ensure acquisition in the same FOV as the first visit.

^1^HMRS is performed using single-voxel spectroscopy (SVS) with voxel placement in the DLPFC to quantify glutamatergic and related metabolites. Both unsuppressed water reference and water-suppressed spectra are collected (voxel size = 10 × 40 × 15 mm³; TR = 3000 ms; TE = 23 ms; 64 averages for water-suppressed spectra and 8 averages for water reference; spectral bandwidth = 2000 Hz; vector size = 2048). Water-suppressed spectra are acquired with RF-based water suppression (bandwidth = 150 Hz) and four preparation scans prior to acquisition. A reference voltage calibration scan is also collected (TR = 12000 ms; TE = 24 ms; 6 measurements) to determine transmit amplitude before spectroscopy acquisition.

### Neuroimaging Processing

As part of the PMADS study protocol, all imaging data will be processed using open-source pipelines to ensure reproducibility and harmonization across modalities. Structural images (T1-weighted, and T2-weighted when available) will be processed with sMRIPrep. Functional MRI data will be minimally preprocessed with fMRIPrep, followed by post-processing and denoising with ICA-AROMA, tedana, and XCP-D to enable generation of PFNs. Diffusion-weighted images will be processed with QSIPrep and reconstructed with QSIRecon. ASL images will be processed with ASLPrep, leveraging outputs from structural preprocessing for co-registration and normalization. Additional quality control will be performed using MRIQC and manual inspection to ensure data integrity across modalities.

#### Structural image processing

T1-weighted structural images will be processed using sMRIPrep (built within the fMRIPrep framework). The pipeline will correct for intensity non-uniformity with N4BiasFieldCorrection, apply skull-stripping, and perform brain tissue segmentation using FSL FAST. Brain surfaces will be reconstructed with FreeSurfer, and pial surfaces will be refined with the T2-weighted image when available. Images will then be spatially normalized to standard templates (MNI152NLin6Asym and MNI152NLin2009cAsym) using nonlinear registration implemented in ANTs. Output will include both volumetric (NIfTI) and surface-based (CIFTI) representations resampled onto the fsLR template. Quality control will be performed using MRIQC and manual visual inspection to identify artifacts or preprocessing failures.

#### Multi-echo fMRI

Resting-state functional MRI data will be preprocessed using fMRIPrep, which will include motion correction, susceptibility distortion correction with TOPUP, co-registration to structural images, and spatial normalization to MNI152NLin6Asym space. Multi-echo time series will then be denoised with tedana (94), which will decompose signals into BOLD-related and non-BOLD components to improve sensitivity and reduce motion and physiological noise. Independent component analysis with ICA-AROMA will additionally be applied to further classify and remove noise components. Post-processed fMRI data will be passed to XCP-D, which will implement nuisance regression (including physiological regressors when available), band-pass filtering, and surface-based resampling onto the fsLR template. The resulting time series will be used to generate PFNs including individualized parcellation and network identification (23,24,26,95). All preprocessing and denoising steps will be accompanied by automated reports and manual quality checks to ensure robust data suitable for downstream PFN analyses.

#### dMRI Processing

Diffusion-weighted images will be preprocessed using QSIPrep, which will include correction for susceptibility distortions, motion, and eddy currents, along with intensity normalization. The pipeline will generate preprocessed diffusion series in standard space, along with confound regressors for motion and signal quality. Following QSIPrep, diffusion reconstruction will be performed using QSIRecon, which will apply generalized q-sampling imaging (GQI) to estimate orientation distribution functions (ODFs) and generate tractography outputs. Automatic tractography workflows will produce whole-brain connectivity matrices, and bundle statistics will be calculated for major white matter pathways. Quality control will also be implemented at multiple stages, including visual inspection of preprocessed images and assessment of motion and signal-to-noise metrics.

#### ASL Processing

ASL images will be preprocessed using ASLPrep, which builds on structural outputs from sMRIPrep for co-registration and normalization. The pipeline will include motion correction for label and control volumes, susceptibility distortion correction, and generation of calibrated cerebral blood flow (CBF) maps using a single-compartment general kinetic model. Calibration (M0) scans will be incorporated to account for baseline magnetization, with volumes smoothed and averaged prior to CBF quantification.

Quality evaluation metrics will be generated for each CBF map, including framewise displacement, DVARS, and a quality evaluation index (QEI) that assesses similarity of perfusion maps to anatomical reference. Parcellated CBF estimates will be extracted for multiple atlases (e.g., Schaefer, Glasser, Tian subcortical, HCP CIFTI subcortical), enabling harmonized analyses across participants. Manual quality control will supplement automated metrics to ensure robust quantification of perfusion measures.

#### GluCEST Processing

GluCEST data will be processed using established ultra-high-field pipelines described in prior work (96–98). Preprocessing will include correction for B₀ and B₁ field inhomogeneities using acquired field maps and WASSR. GluCEST contrast maps will be computed voxelwise as the asymmetry in magnetization transfer ratio at ±3.0 ppm relative to water, yielding percent GluCEST contrast values indexing local glutamate-related signals. Quality control will include exclusion of voxels with excessive B₀ offsets or inadequate B₁ correction and masking of cerebrospinal fluid and non-brain tissue. GluCEST images will be coregistered to high-resolution MP2RAGE structural images, normalized to standard space, and summarized as regional averages within predefined cortical and subcortical regions of interest.

#### ^1^HMRS Processing

Raw spectral data will be exported in RDA format and processed using Osprey (v2.9.0; Osprey Project, Johns Hopkins University) (99), implemented in MATLAB (R2023b; MathWorks, Natick, MA, USA). Spectra will undergo standard preprocessing, including coil combination, eddy-current correction, frequency and phase alignment, water removal, and frequency referencing. Spectral modeling and quantification will be performed using LCModel (v6.3-1N) implemented within Osprey, with basis sets generated for 7T acquisitions and sequence-specific parameters. Default macromolecular and lipid components provided by Osprey will be included in spectral fitting.

Metabolite estimates will include glutamate and glutamine, as well as additional metabolites such as γ-aminobutyric acid (GABA), N-acetylaspartate, creatine, choline-containing compounds, and myo-inositol where signal quality permits. Spectroscopy voxels will be co-registered to each participant’s high-resolution MP2RAGE structural image, and voxel tissue composition (gray matter, white matter, CSF) will be estimated to enable tissue correction of metabolite concentrations. Tissue- and relaxation-corrected metabolite concentrations will be computed in ‘OspreyQuantify’ following established methods (100).

Quality control will include visual inspection of spectral fits and exclusion of metabolite estimates with high uncertainty (Cramér–Rao lower bound > 20%). Signal-to-noise ratio (SNR) and full-width at half-maximum (FWHM) will also be evaluated. Final metabolite measures will be used to complement GluCEST imaging by providing localized neurochemical estimates within the DLPFC.

## Discussion

This study will yield a longitudinal dataset designed to test pre-specified hypotheses while also supporting future hypothesis generation, advancing the study of PMADs through integration of multimodal neuroimaging, biofluid sampling, and deep clinical phenotyping positioned for open-access use. This cohort design incorporates several key innovations. First, it establishes one of the earliest longitudinal, multimodal neuroimaging cohorts spanning preconception through postpartum that also links two reproductive stages (menstrual cycle and pregnancy) within the same individuals. By anchoring perinatal brain changes to preconception features and menstrual cycle sensitivity, this design will enable dissociation of pre-existing vulnerability from pregnancy-induced neuroplasticity, a distinction that is critical for identifying underlying risk markers rather than correlates of pregnancy itself. Second, the cohort specifically enrolls participants at heightened risk for PMADs and is well-suited to capture clinically meaningful heterogeneity in perinatal affective presentations. Third, the integration of advanced multimodal imaging enables the joint probing of structural, functional, and neurochemical context, providing a mechanistic lens on reproductive brain plasticity. Fourth, it leverages multi-echo fMRI, which will allow for precision functional mapping that captures person-specific features of functional network architecture. This individualized approach is particularly important in the context of pregnancy-related neuroplasticity, where assuming fixed or group-average network anatomy may obscure clinically relevant variation. Collectively, this study is designed to support discovery of mechanistic biomarkers of risk and resilience for PMADs that, with expansion to larger samples, could ultimately inform preconception counseling, individualized risk stratification, and nonpharmacologic intervention development.

The present cohort also establishes a framework for future expansion. Subsequent extensions of this design may include the recruitment of comparison groups, such as individuals without risk factors for PMADs, nulliparous controls, non-gestational partners (to capture shared environmental exposures), and gestational carriers or surrogates (to isolate postpartum caregiving effects), to disentangle pregnancy-specific neurobiological changes from general longitudinal or affect-related effects. Such comparisons will be essential for isolating normative versus risk-related patterns of brain remodeling across reproductive transitions.

Implementing a longitudinal protocol of this scope inevitably involves operational challenges. Recruitment and retention are particularly demanding, especially for participants enrolled prior to conception and followed across pregnancy and postpartum. Flexible scheduling, clinic integration, and sustained participant engagement are therefore central components of the cohort infrastructure. MRI acquisition during pregnancy also requires careful attention to safety, participant comfort, and motion mitigation, while the integration of multimodal assessments necessitates strategies to minimize participant burden and maintain data quality. These considerations reflect both the practical demands and the scientific value of a cohort design that seeks to capture the complexity of reproductive brain plasticity in relation to psychiatric risk. By addressing challenges, this study not only offers a feasible framework for advancing mechanistic understanding of PMADs and informing future approaches to clinical care, but also opens horizons for creating a clinically grounded dataset. For example, beyond biomarker discovery, characterizing how structural and functional brain reorganization unfolds across the perinatal transition, this study may inform the development of targeted, personalized, and nonpharmacologic interventions, including network-informed transcranial magnetic stimulation (TMS). Together, this cohort provides a resource for investigating a wide range of questions in perinatal psychopathology, including the neural correlates of PMADs; the role of obstetric and reproductive health factors in shaping psychiatric outcomes; and the transdiagnostic mechanisms that cut across mood, stress, and caregiving systems during pregnancy and postpartum.

## Supporting information

Supplementary Material 2. Interval Depression Inventory

Supplementary Material 3. Interval Anxiety Inventory

## List of abbreviations

PMADs: Perinatal mood and anxiety disorders
OCP: Oral contraceptive pill
PMDD: Premenstrual dysphoric disorder
MRI: Magnetic resonance imaging
fMRI: Functional magnetic resonance imaging
ASL: Arterial spin labeling
PSST: Premenstrual Symptoms Screening Tool
PTSD: Post-traumatic stress disorder
BPD: Borderline personality disorder
THC: Tetrahydrocannabinol
BMI: Body mass index
DRSP: Daily Record of Severity of Problems
EPDS: Edinburgh Postnatal Depression Scale
PHQ-9: Patient Health Questionnaire-9
BDI-II: Beck Depression Inventory-II
GAD-7: Generalized Anxiety Disorder-7
BAI: Beck Anxiety Inventory
STAI: State-Trait Anxiety Inventory
PrAS: Pregnancy-Related Anxiety Scale
MDQ: Mood Disorder Questionnaire
PSS-10: Perceived Stress Scale-10
OCI: Obsessive–Compulsive Inventory
PSQI: Pittsburgh Sleep Quality Index
BADDS: Brown Attention-Deficit Disorder Scale
PRIME: Psychosis-Risk Screening Questionnaire
DERS-16: Difficulties in Emotion Regulation Scale-16
PCL-5: PTSD Checklist for DSM-5
SLESQ-R: Stressful Life Events Screening Questionnaire-Revised
ACEs: Adverse Childhood Experiences
CTQ-SF: Childhood Trauma Questionnaire-Short Form
BCEs: Benevolent Childhood Experiences
ASQ: Adult/Adolescent Social Support Questionnaire
AHC-HRSN: Accountable Health Communities Health-Related Social Needs Screening Tool
FertiQoL: Fertility Quality of Life Scale
TLFB: Timeline Follow-Back
MAAS: Maternal Antenatal Attachment Scale
MPAS: Maternal Postnatal Attachment Scale
PBQ: Postpartum Bonding Questionnaire
BIMF: Barkin Index of Maternal Functioning
City-BiTS: City Birth Trauma Scale
PSI: Pregnancy Symptoms Inventory
NVP: Nausea and vomiting of pregnancy
PUQE-24: Pregnancy-Unique Quantification of Emesis and Nausea-24
HELP: HyperEmesis Level Prediction
EMR: Electronic medical record
CAT-GOASSESS: Computerized Adaptive Testing version of the GOASSESS
CNB: Computerized Neurocognitive Battery
PVT: Psychomotor Vigilance Test
sPCPT: Short Penn Continuous Performance Test
SLNB2: Short Letter-N-Back (2-back)
AIM: Penn Abstraction, Inhibition, and Working Memory Task
ER40: Penn Emotion Recognition Test (40-item)
ADT: Age Differentiation Test
MEDF: Measured Emotion Differentiation Test
PMAT: Penn Matrix Reasoning Test
PLOT: Penn Line Orientation Test
SPVRT: Short Penn Verbal Reasoning Test
EDTA: Ethylenediamine tetraacetic acid
CBC: Complete blood count
LLT: Left lateral tilt
IVC: Inferior vena cava
nNav: Volumetric navigators
EPI: Echo-planar imaging
pCASL: Pseudo-continuous arterial spin labeling
¹H-MRS: Proton magnetic resonance spectroscopy
DLPFC: Dorsolateral prefrontal cortex
PFN: Personalized functional network
GQI: Generalized q-sampling imaging
ODF: Orientation distribution function
CBF: Cerebral blood flow
QEI: Quality evaluation index
GABA: γ-aminobutyric acid
SNR: Signal-to-noise ratio
FWHM: full-width at half-maximum
WASSR: Water saturation shift referencing
TMS: Transcranial magnetic stimulation

## Declarations

### Ethics approval and consent to participate

This study was approved by the Institutional Review Board (IRB) of the University of Pennsylvania (protocol number: 856163). All participants provided written informed consent prior to participation.

### Consent for publication

Not applicable

### Availability of data and materials

Data sharing is not applicable to this article as no datasets were generated or analyzed during the current stage of the study. Upon completion of data collection and primary analyses, de-identified data generated will be made publicly available via an open-access repository, in accordance with IRB guidelines and participant consent procedures. Access to certain data elements may be restricted to protect participant privacy.

No results, data, or figures in this manuscript have been published elsewhere, nor are they under consideration by another publisher.

### Authors’ contributions

S.S. conceived the study. N.R.-A., C.H., A.F., S.R., L.P., and S.S. designed the overall research framework and developed the study protocol. D.H.W., D.R.R., M.E.C., S.S., L.H., E.B.B., R.B., R.K., T.M.M., S.L.K., J.C.S., L.K.W., and D.I. contributed to the design of the clinical assessments, study-specific questionnaires, cognitive assessments, specimen processing procedures, and recruitment procedures used and their integration into the study protocol clinical. D.R.R., T.S., M.C., M.A.E., H.H., M.T., M.D.T., C.B.K., J.A.D., and A.V. contributed to the design of neuroimaging procedures including acquisition protocols and processing pipelines. N.R.-A., C.H., A.F., S.R., H.R., M.I., L.Hu., and S.S. contributed to the implementation of the study, including participant recruitment strategies, study coordination, and data collection procedures. N.R.-A. and S.S. drafted the manuscript. L.P., C.H., E.M.B., R.B., M.E.C., S.L.K., J.C.S., T.S., M.C., A.S.K., L.Hu., K.Z., S.S., L.H., D.I., S.D., R.K., L.K.W., M.A.E., H.H., M.T., M.D.T., C.B.K., J.A.D., A.V., T.M.M., D.H.W., and D.R.R. substantially revised the manuscript and provided critical feedback. All authors approved the submitted version of the manuscript.

## Acknowledgements

We thank the Center for Advanced Magnetic Resonance Imaging and Spectroscopy at the University of Pennsylvania (RRID: SCR_022398) for their assistance with the acquisition of MRI data.

