## Supplemental Data 1 for "The PMADS Project: A Longitudinal Multimodal Cohort Study to Understand Risk for Perinatal Mood and Anxiety Disorders"

#### Reproductive Mental Health History Questionnaire (G0 Version)

*Personal History*

**1.** Have you ever been diagnosed with any of the following conditions? (Check all that apply.)

☐ Premenstrual dysphoric disorder (PMDD)
☐ Premenstrual syndrome (PMS)*
☐ Premenstrual exacerbation of an underlying mood disorder (PME)
☐ Borderline personality disorder (BPD)
☐ Bipolar disorder
☐ Major Depression Disorder (MDD)
☐ Generalized Anxiety Disorder (GAD)
☐ Obsessive-Compulsive Disorder (OCD)
☐ Post-Traumatic Stress Disorder (PTSD)

** In this study, premenstrual syndrome (PMS) refers to a clinical condition involving a pattern of recurrent physical and/or emotional symptoms that occur during the 5 days before menstruation and interfere with daily functioning. This does not refer to mild emotional or physical symptoms that occur 1-2 days before menses but that do not interfere with daily functioning..*

---

*If you have received a diagnosis of Premenstrual dysphoric disorder (PMDD):*

**A1**. At what age did your symptoms of *PMDD* start? ________ years old
**A2**. At what age were you diagnosed with *PMDD*? ________ years old
**A3**. Who diagnosed you with *PMDD*? (Check one.)

☐ Psychiatrist
☐ Mental health provider other than a psychiatrist
☐ Primary care provider
☐ OB/GYN
☐ Other (please specify): ______________________________

*If diagnosed by a psychiatrist or other mental health provider:* Does the provider who diagnosed you with *PMDD* specialize in women's mental health (e.g. reproductive psychiatrist, perinatal psychiatrist, reproductive psychologist, etc.)?
☐ Yes ☐ No ☐ I don't know

**A4.** Before you were diagnosed with *PMDD*, did you complete daily symptom ratings for at least 2 months (for example, the Daily Record of Severity of Problems or similar daily tracking form)?

☐ Yes
☐ No
☐ Not sure

**A5.** Were those daily symptom ratings reviewed by the provider who diagnosed you with *PMDD* as part of making the diagnosis?

☐ Yes
☐ No
☐ Not sure

---

**A5**. Are you currently being treated for *PMDD*?

☐ Yes ☐ No

**A6**. Are you currently taking medication for *PMDD*?

☐ Yes ☐ No

---

*If you have received a diagnosis of Premenstrual syndrome (PMS)*:*

** In this study, premenstrual syndrome (PMS) refers to a clinical condition involving a pattern of recurrent physical and/or emotional symptoms that occur during the 5 days before menstruation and interfere with daily functioning. This does not refer to mild emotional or physical symptoms that occur 1-2 days before menses but that do not interfere with daily functioning..*

**A1**. At what age did your symptoms of *PMS* start? ________ years old

**A2**. At what age were you diagnosed with *PMS*? ________ years old

**A3**. Who diagnosed you with *PMS*? (Check one.)

☐ Psychiatrist

☐ Mental health provider other than a psychiatrist

☐ Primary care provider

☐ OB/GYN

☐ Other (please specify): ______________________________

*If diagnosed by a psychiatrist or other mental health provider:* Does the provider who diagnosed you with *PMS* specialize in women's mental health (e.g. reproductive psychiatrist, perinatal psychiatrist, reproductive psychologist, etc.)?

☐ Yes ☐ No ☐ I don't know

**A4**. Are you currently being treated for *PMS*?

☐ Yes ☐ No

**A5**. Are you currently taking medication for *PMS*?

☐ Yes ☐ No

---

*If Premenstrual exacerbation of an underlying mood disorder (PME):*

**A1**. At what age did your symptoms of *PME* start? ________ years old

For which diagnosed condition(s) do your symptoms worsen before your menstrual period? (Check all that apply.)

☐ Major depressive disorder

☐ Generalized anxiety disorder

☐ Bipolar disorder

☐ Post-traumatic stress disorder (PTSD)

☐ Panic disorder

☐ Obsessive-compulsive disorder (OCD)

☐ Other (please specify): ______________________________

**A2**. At what age were you diagnosed with *PME*? ________ years old

**A3**. Who diagnosed you with *PME*? (Check one.)

---

☐ Psychiatrist
☐ Mental health provider other than a psychiatrist
☐ Primary care provider
☐ OB/GYN
☐ Other (please specify): ______________________________

*If diagnosed by a psychiatrist or other mental health provider:* Does the provider who diagnosed you with *PME* specialize in women's mental health (e.g. reproductive psychiatrist, perinatal psychiatrist, reproductive psychologist, etc.)?

☐ Yes ☐ No ☐ I don't know

**A4.** Before you were diagnosed with *PME*, did you complete daily symptom ratings for at least 2 months (for example, the Daily Record of Severity of Problems or similar daily tracking form)?

☐ Yes
☐ No
☐ Not sure

**A5.** Were those daily symptom ratings reviewed by the provider who diagnosed you with *PME* as part of making the diagnosis?

☐ Yes
☐ No
☐ Not sure

**A6**. Are you currently being treated for *PME*?

☐ Yes ☐ No

**A7**. Are you currently taking medication for *PME*?

☐ Yes ☐ No

---

*If you have received a diagnosis of Borderline personality disorder (BPD):*

**A1**. At what age did your symptoms of *BPD* start? ________ years old

**A2**. At what age were you diagnosed with *BPD*? ________ years old

**A3**. Who diagnosed you with *BPD*? (Check one.)

☐ Psychiatrist
☐ Mental health provider other than a psychiatrist
☐ Primary care provider
☐ OB/GYN
☐ Other (please specify): ______________________________

*If diagnosed by a psychiatrist or other mental health provider:* Does the provider who diagnosed you with *BPD* specialize in women's mental health (e.g. reproductive psychiatrist, perinatal psychiatrist, reproductive psychologist, etc.)?

☐ Yes ☐ No ☐ I don't know

**A4**. Are you currently being treated for *BPD*?

☐ Yes ☐ No

**A5**. Are you currently taking medication for *BPD*?

☐ Yes ☐ No

---

---

*If you have received a diagnosis of Bipolar Disorder:*

**A1**. At what age did your symptoms of *Bipolar Disorder* start? ________ years old

**A2**. At what age were you diagnosed with *Bipolar Disorder*? ________ years old

**A3**. Who diagnosed you with *Bipolar Disorder*? (Check one.)

☐ Psychiatrist

☐ Mental health provider other than a psychiatrist

☐ Primary care provider

☐ OB/GYN

☐ Other (please specify): ______________________________

*If diagnosed by a psychiatrist or other mental health provider:* Does the provider who diagnosed you with *Bipolar Disorder* specialize in women's mental health (e.g. reproductive psychiatrist, perinatal psychiatrist, reproductive psychologist, etc.)?

☐ Yes ☐ No ☐ I don't know

**A4**. Are you currently being treated for *Bipolar Disorder*?

☐ Yes ☐ No

**A5**. Are you currently taking medication for *Bipolar Disorder*?

☐ Yes ☐ No

---

**2.** Do you experience worsening mood symptoms (such as anger, irritability, anxiety, or depression) in the week before your period that improves when your period starts? Only consider cycles where you were <u>not</u> using oral contraceptive pills, transdermal contraceptive patch, contraceptive shot, contraceptive ring, had an arm implant, or had a hormonal IUD without a period.

☐ Yes

☐ No

☐ Not currently, but I have in the past

***If Yes, continue to B1.***

***If Not currently, but I have in the past, skip to B2.***

***If No, skip to Question 3***

**B1.** Do you experience any of these symptoms in the week before your period? Only consider cycles where you were <u>not</u> using oral contraceptive pills, transdermal contraceptive patch, contraceptive shot, contraceptive ring, had an arm implant, or had a hormonal IUD without a period.

*Check all that apply*

☐ Mood swings (suddenly feeling sad or tearful)

☐ Feeling unusually sensitive to criticism or rejection

☐ Feeling especially irritable or angry

☐ Frequent arguments or conflicts with others

☐ Feeling very sad, down, depressed, or hopeless

☐ Feeling overly self-critical
☐ Feeling extremely anxious, tense, or on edge
☐ Loss of interest in work, school, friends, or hobbies
☐ Difficulty concentrating
☐ Very low energy or fatigue
☐ Increased appetite or specific food cravings
☐ Sleeping more than usual or difficulty sleeping
☐ Feeling overwhelmed or out of control
☐ Physical symptoms (e.g., breast tenderness, bloating, joint/muscle pain, weight gain)

*[If checks any of the above]*
**B1a.** What happens to these symptoms after your period starts:
☐ The symptoms start to improve within a few days after the onset of menses and become minimal or absent in the week after menses
☐ The symptoms start to improve within a few days after the onset of menses and remain present at a lower level the remainder of the cycle
☐ The symptoms get worse with the onset of menses and improve when menses ends
☐ The symptoms stay the same; there is not a relationship between menses and mood symptoms.
☐ Other (please specify): ____________________________________________

**B1b.** Over the last 12 cycles, how often did you experience these symptoms? Only consider cycles where you were not using oral contraceptive pills, transdermal contraceptive patch, contraceptive shot, contraceptive ring, had an arm implant, or had a hormonal IUD without a period.
☐ Every cycle
☐ Almost every cycle (10-11 cycles)
☐ More often than not (6-9 cycles)
☐ Less than half of cycles (1-5 cycles)
☐ Don't know

**B2.** When did you last experience these symptoms? ________________________________

**B2a.** Think about the time when you experienced these symptoms. On average over the course of 12 cycles, how often did you experience these symptoms? Only consider cycles where you were not using oral contraceptive pills, transdermal contraceptive patch, contraceptive shot, contraceptive ring, had an arm implant, or had a hormonal IUD without a period.

☐ Every cycle
☐ Almost every cycle (10-11 cycles)
☐ More often than not (6-9 cycles)
☐ Less than half of cycles (1-5 cycles)
☐ Don't know

**B2b.**What, if anything, made these symptoms improve?

- ☐ Hormonal contraception
- ☐ Psychiatric medication
- ☐ Psychotherapy
- ☐ Other (please specify): ______________________________________________

**B3.** To what extent have these symptoms affected your relationships or your interactions with other people? For example, have they caused you any problems in your relationships with your family, romantic partner or friends?

- ☐ Not at all
- ☐ Mild
- ☐ Moderate
- ☐ Severe

**B4.** To what extent have these symptoms affected your work/school? For example, have they affected your attendance at work or school? Have they affected the quality of your work/schoolwork?

- ☐ Not at all
- ☐ Mild
- ☐ Moderate
- ☐ Severe

**B5.** To what extent have these symptoms affected your ability to take care of things at home? For example, have they affected your ability to do other things that are important to you like religious activities, physical exercise, or hobbies? Did you avoid doing anything because you felt like you weren't up to it?

- ☐ Not at all
- ☐ Mild
- ☐ Moderate
- ☐ Severe

**B6.** How much have you been bothered or upset by having these symptoms?

- ☐ Not at all
- ☐ Mild
- ☐ Moderate
- ☐ Severe

**3.** Have you ever experienced new or worsening mood symptoms such as anger, irritability, anxiety, or depression in response to taking any of the following medications?
*Check all that apply.*

- ☐ Oral contraceptive pills (e.g birth control pills)
- ☐ Transdermal contraceptive patch (e.g. birth control patch)
- ☐ Arm implant (e.g. Nexplanon, Implanon, etonogestrel implant)
- ☐ Hormonal intrauterine device (IUD; e.g. Mirena, Lyletta, Skyla, Kyleena)
- ☐ Injectable contraceptive (e.g. birth control shot, DepoProvera)

☐ Contraceptive vaginal ring (e.g. birth control ring, Nuvaring)
☐ Fertility treatments (e.g. Clomid, Letrozole, Ovidrel, hCG, FSH, etc)
☐ Other hormonal medications (please specify) ___________________________________

---

*If Oral contraceptive pills (e.g birth control pills):*

**C1.** Which of the following symptoms started or became worse after starting oral contraceptive pills? How severe were these symptoms when you were taking oral contraceptive pills? If you have experienced mood symptoms in response to more than one type/brand of oral contraceptive pill, respond below thinking of the type/brand of oral contraceptive pill that caused the most severe or most consistent mood symptoms.

| **Symptom** | **Not at all** | **Mild** | **Moderate** | **Severe** |
|---|---|---|---|---|
| 1. Anger/irritability | | | | |
| 2. Anxiety/tension | | | | |
| 3. Tearful/increased sensitivity to rejection | | | | |
| 4. Depressed mood/hopelessness | | | | |
| 5. Decreased interest in work activities | | | | |
| 6. Decreased interest in home activities | | | | |
| 7. Decreased interest in social activities | | | | |
| 8. Difficulty concentrating | | | | |
| 9. Fatigue/lack of energy | | | | |
| 10. Overeating/food cravings | | | | |
| 11. Insomnia | | | | |
| 12. Hypersomnia (needing more sleep) | | | | |
| 13. Feeling overwhelmed or out of control | | | | |
| 14. Physical symptoms: breast tenderness, headaches, joint/muscle pain, bloating, weight gain | | | | |

**C2.** Did the symptoms you experienced interfere with:

| **Symptom** | **Not at all** | **Mild** | **Moderate** | **Severe** |
|---|---|---|---|---|
| A. Your work efficiency or productivity | | | | |
| B. Your relationships with coworkers | | | | |
| C. Your relationships with your family | | | | |

| |
|---|
| D. Your social life activities |
| E. Your home responsibilities |

**C3.** From the options below, select the schematic that most looks like the oral contraceptive pill that led to these symptoms. Colors in the images may be different from colors in the pill pack you used. Focus primarily on the number of different pill colors and number of each pill of a given color. *If possible, please check your prescription history, pill packaging, or medication records before answering.*

☐ Monophasic

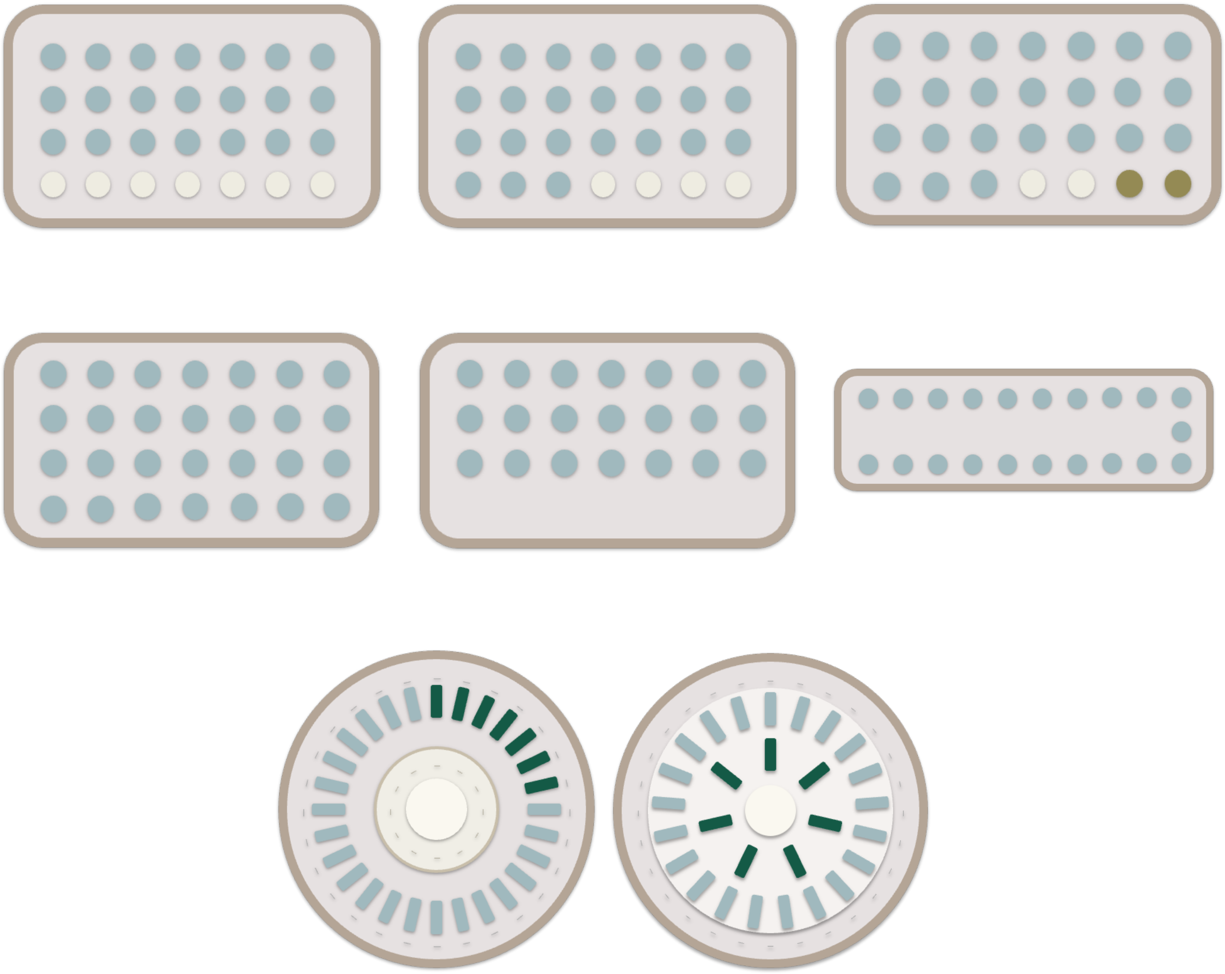

☐ Biphasic

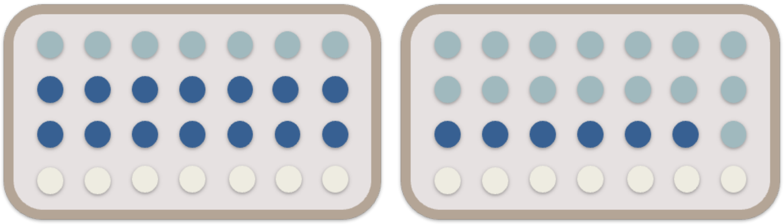

- ☐ Triphasic

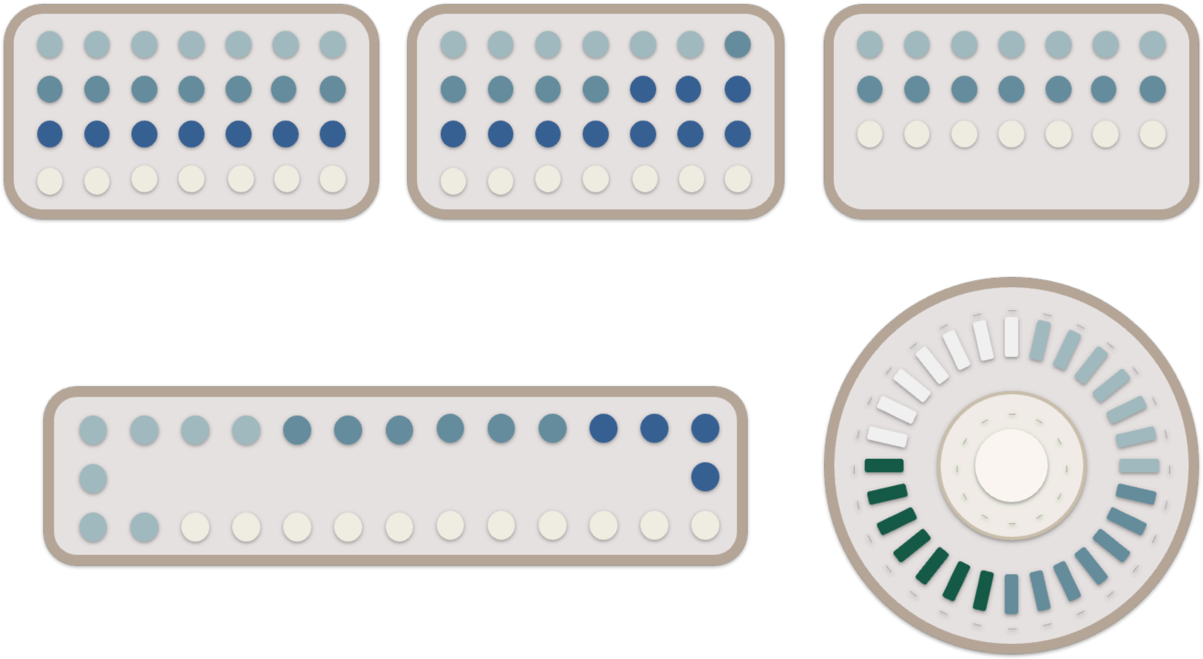

- ☐ I don’t know
- ☐ Other

*[If any response except I don’t know]*

How confident are you in your answer about which oral contraceptive pill led to these symptoms?

- ☐ I confirmed this using medical records, prescription history, or packaging
- ☐ Very confident — I clearly remember the exact pill or brand
- ☐ Moderately confident — I think this is correct but am not completely sure
- ☐ Slightly confident — this is my best guess
- ☐ Not confident — I am unsure which pill it was

**C4.** If you know which oral contraceptive pill led to these symptoms, please provide it here. If you have experienced mood symptoms in response to more than one type/brand of oral contraceptive pill, please provide the type/brand of oral contraceptive pill that caused the most severe or most consistent mood symptoms. *If possible, please check your prescription history, pill packaging, or medication records before answering.*

☐ Yes

______________________________________________________________

☐ I don't know

*[If any response except I don't know]*

How confident are you in your answer about which oral contraceptive pill led to these symptoms?

☐ I confirmed this using medical records, prescription history, or packaging
☐ Very confident — I clearly remember the exact pill or brand
☐ Moderately confident — I think this is correct but am not completely sure
☐ Slightly confident — this is my best guess
☐ Not confident — I am unsure which pill it was

**C5.** When did you start taking this oral contraceptive pill? Please provide the month and year if possible. Some people find it easier to recall this information if thinking about how old they were when they started taking it. ________________________________

**C6.** How confident are you in your answer about when you started taking this oral contraceptive pill?

☐ I confirmed this using medical records, prescription history, or another dated record
☐ Very confident — I clearly remember the month and year
☐ Moderately confident — I think the month and year are correct, but I am not completely sure
☐ Slightly confident — this is my best guess
☐ Not confident — I am unsure when I started taking it

**C7.** For how long did you take this type of oral contraceptive pill?

☐ Less than 1 week
☐ 1 week - 1 month
☐ 1 - 3 months
☐ 3 - 6 months
☐ 6 months or more
☐ Unsure

**C8.** How long after starting this oral contraceptive pill did you first notice new or worsening mood symptoms?

☐ The day I started the pill
☐ Within 1 week after starting the pill
☐ Within 1 month after starting the pill
☐ Within 3 months after starting the pill
☐ Within 6 months after starting the pill

☐ Longer than 6 months after starting the pill
☐ Unsure
☐ Other (please specify) ____________________________________________

**C9.** For how long did you experience the new or worsening mood symptoms while taking this oral contraceptive pill?
☐ 1 week or less
☐ 1 week - 1 month
☐ 1 - 3 months
☐ 3 - 6 months
☐ 6 months or more
☐ Unsure
☐ Other (please specify) ____________________________________________

**C10.** How long did you continue taking this oral contraceptive pill after noticing the new or worsening mood symptoms?
☐ 1 week or less
☐ 1 week - 1 month
☐ 1 - 3 months
☐ 3 - 6 months
☐ 6 months or more
☐ Unsure
☐ Other (please specify) ____________________________________________

**C11.** Did the new/worsening mood symptoms you experienced when starting this oral contraceptive pill go away when you stopped taking this medication?
◻ Yes, completely
◻ Yes, partially
◻ No
◻ Not Applicable (please specify) ______________________________________

*[If yes]*
After you stopped taking this oral contraceptive pill, how long did it take for the new/worsening mood symptoms to improve or go away?
◻ Within a few days after stopping the pill
◻ Within 1 week after stopping the pill
◻ Within 1 month after stopping the pill
◻ Within 3 months after stopping the pill
◻ Longer than 3 months after stopping the pill
◻ Unsure
◻ Other (please specify) ________________________________________

**C12.** Did you experience the above mood symptoms in response to more than one type/brand of oral contraceptive pill?

- ☐ Yes
- ☐ No

*If Transdermal Contraceptive Patch (e.g. birth control patch):*

**C1.** Which of the following symptoms started or became worse after starting the transdermal contraceptive patch? How severe were these symptoms when you were using the patch? If you have experienced mood symptoms in response to more than one type/brand of transdermal contraceptive patch, respond below thinking of the type/brand of transdermal contraceptive patch that caused the most severe or most consistent mood symptoms.

| **Symptom** | **Not at all** | **Mild** | **Moderate** | **Severe** |
|---|---|---|---|---|
| 1. Anger/irritability | | | | |
| 2. Anxiety/tension | | | | |
| 3. Tearful/increased sensitivity to rejection | | | | |
| 4. Depressed mood/hopelessness | | | | |
| 5. Decreased interest in work activities | | | | |
| 6. Decreased interest in home activities | | | | |
| 7. Decreased interest in social activities | | | | |
| 8. Difficulty concentrating | | | | |
| 9. Fatigue/lack of energy | | | | |
| 10. Overeating/food cravings | | | | |
| 11. Insomnia | | | | |
| 12. Hypersomnia (needing more sleep) | | | | |
| 13. Feeling overwhelmed or out of control | | | | |
| 14. Physical symptoms: breast tenderness, headaches, joint/muscle pain, bloating, weight gain | | | | |

**C2.** Did the symptoms you experienced interfere with:

| **Symptom** | **Not at all** | **Mild** | **Moderate** | **Severe** |
|---|---|---|---|---|
| A. Your work efficiency or productivity | | | | |
| B. Your relationships with coworkers | | | | |

| |
|---|
| C. Your relationships with your family |
| D. Your social life activities |
| E. Your home responsibilities |

**C3.** If you know which transdermal contraceptive patch led to these symptoms, please provide it here. If you have experienced mood symptoms in response to more than one type/brand of transdermal contraceptive patch, please provide the type/brand of the patch that caused the most severe or most consistent mood symptoms. *If possible, please check your prescription history, packaging, or medication records before answering.*

☐ Yes

____________________________________________________________

☐ I don't know

*[If any response except I don't know]*

How confident are you in your answer about which transdermal contraceptive patch led to these symptoms?

☐ I confirmed this using medical records, prescription history, or packaging
☐ Very confident — I clearly remember the exact patch or brand
☐ Moderately confident — I think this is correct but am not completely sure
☐ Slightly confident — this is my best guess
☐ Not confident — I am unsure which patch it was

**C4.** When did you start using this transdermal contraceptive patch? Please provide the month and year if possible. Some people find it easier  to recall this information if thinking about how old they were when they started using it. __________________________________

**C5.** How confident are you in your answer about when you started using this transdermal contraceptive patch?

☐ I confirmed this using medical records, prescription history, or another dated record
☐ Very confident — I clearly remember the month and year
☐ Moderately confident — I think the month and year are correct, but I am not completely sure
☐ Slightly confident — this is my best guess
☐ Not confident — I am unsure when I started taking it

**C6.** For how long did you use this type of transdermal contraceptive patch?

☐ Less than 1 week
☐ 1 week - 1 month
☐ 1 - 3 months
☐ 3 - 6  months
☐ 6 months or more
☐ Unsure

**C7.** How long after starting this transdermal contraceptive patch did you first notice new or worsening mood symptoms?

- ◻ The day I started the patch
- ◻ Within 1 week after starting the patch
- ◻ Within 1 month after starting the patch
- ◻ Within 3 months after starting the patch
- ◻ Within 6 months after starting the patch
- ◻ Longer than 6 months after starting the patch
- ◻ Unsure
- ◻ Other (please specify) ________________________________________________

**C8.** For how long did you experience the new or worsening mood symptoms while using this transdermal contraceptive patch?

- ◻ 1 week or less
- ◻ 1 week - 1 month
- ◻ 1 - 3 months
- ◻ 3 - 6 months
- ◻ 6 months or more
- ◻ Unsure
- ◻ Other (please specify) ________________________________________________

**C9.** For how long did you continue using the patch after noticing the new or worsening mood symptoms?

- ☐ Less than 1 week
- ☐ 1 week - 1 month
- ☐ 1 - 3 months
- ☐ 3 - 6 months
- ☐ 6 months or more
- ☐ Unsure
- ☐ Other (please specify) ________________________________________________

**C10.** Did the new/worsening mood symptoms you experienced when starting this transdermal contraceptive patch go away when you stopped using this patch?

- ◻ Yes, completely
- ◻ Yes, partially
- ◻ No
- ◻ Not Applicable (please specify) __________________________________________

  *[If yes]*

  After you stopped using this transdermal contraceptive patch, how long did it take for the new/worsening mood symptoms to improve or go away?

  - ◻ Within a few days after stopping the patch
  - ◻ Within 1 week after stopping the patch

- ☐ Within 1 month after stopping the patch
- ☐ Within 3 months after stopping the patch
- ☐ Within 6 months after starting the patch
- ☐ Longer than 6 months after starting the patch
- ☐ Unsure
- ☐ Other (please specify) ________________________________________________

**C11.** Did you experience the above mood symptoms in response to more than one type/brand of transdermal contraceptive patch?

- ☐ Yes
- ☐ No

---

*If Contraceptive vaginal ring (e.g. birth control ring, Nuvaring):*

**C1.** Which of the following symptoms started or became worse when you were using the contraceptive vaginal ring? How severe were these symptoms when you were using the ring? If you have experienced mood symptoms in response to more than one type/brand of contraceptive vaginal ring, respond below thinking of the type/brand of contraceptive vaginal ring that caused the most severe or most consistent mood symptoms.

| Symptom | Not at all | Mild | Moderate | Severe |
|---|---|---|---|---|
| 1. Anger/irritability | | | | |
| 2. Anxiety/tension | | | | |
| 3. Tearful/increased sensitivity to rejection | | | | |
| 4. Depressed mood/hopelessness | | | | |
| 5. Decreased interest in work activities | | | | |
| 6. Decreased interest in home activities | | | | |
| 7. Decreased interest in social activities | | | | |
| 8. Difficulty concentrating | | | | |
| 9. Fatigue/lack of energy | | | | |
| 10. Overeating/food cravings | | | | |
| 11. Insomnia | | | | |
| 12. Hypersomnia (needing more sleep) | | | | |
| 13. Feeling overwhelmed or out of control | | | | |
| 14. Physical symptoms: breast tenderness, headaches, joint/muscle pain, bloating, weight gain | | | | |

**C2.** Did the symptoms you experienced interfere with:

| **Symptom** | **Not at all** | **Mild** | **Moderate** | **Severe** |
|---|---|---|---|---|
| A. Your work efficiency or productivity | | | | |
| B. Your relationships with coworkers | | | | |
| C. Your relationships with your family | | | | |
| D. Your social life activities | | | | |
| E. Your home responsibilities | | | | |

**C3.** If you know which contraceptive vaginal ring led to these symptoms, please provide it here. If you have experienced mood symptoms in response to more than one type/brand of contraceptive vaginal ring, please provide the type/brand of the ring that caused the most severe or most consistent mood symptoms. *If possible, please check your prescription history, packaging, or medication records before answering.*

☐ Yes

_______________________________________________

☐ I don't know

*[If any response except I don't know]*

How confident are you in your answer about which contraceptive vaginal ring led to these symptoms?

☐ I confirmed this using medical records, prescription history, or packaging
☐ Very confident — I clearly remember the exact ring or brand
☐ Moderately confident — I think this is correct but am not completely sure
☐ Slightly confident — this is my best guess
☐ Not confident — I am unsure which ring it was

**C4.** When did you start using this contraceptive vaginal ring? Please provide the month and year if possible. Some people find it easier to recall this information if thinking about how old they were when they started using it. ________________________________

**C5.** How confident are you in your answer about when you started taking this contraceptive vaginal ring?

☐ I confirmed this using medical records, prescription history, or another dated record
☐ Very confident — I clearly remember the month and year
☐ Moderately confident — I think the month and year are correct, but I am not completely sure
☐ Slightly confident — this is my best guess
☐ Not confident — I am unsure when I started taking it

**C6.** For how long did you use this type of contraceptive vaginal ring?

- ☐ Less than 1 week
- ☐ 1 week - 1 month
- ☐ 1 - 3 months
- ☐ 3 - 6 months
- ☐ 6 months or more
- ☐ Unsure

**C5.** How long after starting this contraceptive vaginal ring did you first notice new or worsening mood symptoms?

- ▫ The day I started the ring
- ▫ Within 1 week after starting the ring
- ▫ Within 1 month after starting the ring
- ▫ Within 3 months after starting the ring
- ▫ Within 6 months after starting the ring
- ▫ Longer than 6 months after starting the ring
- ▫ Unsure
- ▫ Other (please specify) ________________________________________________

**C6.** For how long did you experience the new or worsening mood symptoms while using this contraceptive vaginal ring?

- ☐ 1 week or less
- ☐ 1 week - 1 month
- ☐ 1 - 3 months
- ☐ 3 - 6 months
- ☐ 6 months or more
- ☐ Unsure
- ☐ Other (please specify) ________________________________________________

**C7.** For how long did you continue using this contraceptive vaginal ring after noticing the new or worsening mood symptoms?

- ☐ Less than 1 week
- ☐ 1 week - 1 month
- ☐ 1 - 3 months
- ☐ 3 - 6 months
- ☐ 6 months or more
- ☐ Unsure
- ☐ Other (please specify) ________________________________________________

**C8.** Did the new/worsening mood symptoms you experienced when starting this contraceptive vaginal ring go away when you stopped using this ring?

- ▫ Yes, completely

- ☐ Yes, partially
- ☐ No
- ☐ Not Applicable (please specify) ____________________________________________

  *[If yes]*

  After you stopped taking this contraceptive vaginal ring, how long did it take for the new/worsening mood symptoms to improve or go away?
  - ☐ Within a few days after stopping the ring
  - ☐ Within 1 week after stopping the ring
  - ☐ Within 1 month after stopping the ring
  - ☐ Within 3 months after starting the ring
  - ☐ Within 6 months after starting the ring
  - ☐ Longer than 6 months after starting the ring
  - ☐ Unsure
  - ☐ Other (please specify) ____________________________________________

**C9.** Did you experience the above mood symptoms in response to more than one type/brand of contraceptive vaginal ring?
- ☐ Yes
- ☐ No

---

*If Arm implant (e.g. Nexplanon, Implanon, etonogestrel implant):*

**C1.** Which of the following symptoms started or became worse after the arm implant was inserted? How severe were these symptoms? If you have experienced mood symptoms in response to more than one type/brand of arm implant, respond below thinking of the arm implant that caused the most severe or most consistent mood symptoms.

| **Symptom** | **Not at all** | **Mild** | **Moderate** | **Severe** |
|---|---|---|---|---|
| 1. Anger/irritability | | | | |
| 2. Anxiety/tension | | | | |
| 3. Tearful/increased sensitivity to rejection | | | | |
| 4. Depressed mood/hopelessness | | | | |
| 5. Decreased interest in work activities | | | | |
| 6. Decreased interest in home activities | | | | |
| 7. Decreased interest in social activities | | | | |
| 8. Difficulty concentrating | | | | |
| 9. Fatigue/lack of energy | | | | |
| 10. Overeating/food cravings | | | | |

| |
|---|
| 11. Insomnia |
| 12. Hypersomnia (needing more sleep) |
| 13. Feeling overwhelmed or out of control |
| 14. Physical symptoms: breast tenderness, headaches, joint/muscle pain, bloating, weight gain |

**C2.** Did the symptoms you experienced interfere with:

| **Symptom** | **Not at all** | **Mild** | **Moderate** | **Severe** |
|---|---|---|---|---|
| A. Your work efficiency or productivity | | | | |
| B. Your relationships with coworkers | | | | |
| C. Your relationships with your family | | | | |
| D. Your social life activities | | | | |
| E. Your home responsibilities | | | | |

**C3.** If you know which arm implant led to these symptoms, please provide it here. If you have experienced mood symptoms in response to more than one type/brand of arm implant, please provide the type/brand of the implant that caused the most severe or most consistent mood symptoms. *If possible, please check your prescription history, packaging, or medication records before answering.*

☐ Yes

____________________________________________________________

☐ I don't know

*[If any response except I don't know]*

How confident are you in your answer about which arm implant(s) led to these symptoms?

☐ I confirmed this using medical records, prescription history, or packaging
☐ Very confident — I clearly remember the exact implant or brand
☐ Moderately confident — I think this is correct but am not completely sure
☐ Slightly confident — this is my best guess
☐ Not confident — I am unsure which implant it was

**C4.** When did you start using this arm implant? Please provide the month and year if possible. Some people find it easier to recall this information if thinking about how old they were when they started using it. ______________________________

**C5.** How confident are you in your answer about when you started taking this arm implant?

☐ I confirmed this using medical records, prescription history, or another dated record

☐ Very confident — I clearly remember the month and year
☐ Moderately confident — I think the month and year are correct, but I am not completely sure
☐ Slightly confident — this is my best guess
☐ Not confident — I am unsure when I started taking it

**C6.** For how long did you have the arm implant?

☐ Less than 1 week
☐ 1 week - 1 month
☐ 1 - 3 months
☐ 3 - 6 months
☐ 6 months or more
☐ Unsure

**C7.** How long after the arm implant was inserted did you first notice new or worsening mood symptoms?

▫ Within 1 day
▫ Within 1 week
▫ Within 1 month
▫ Within 3 months
▫ Within 6 months
▫ Longer than 6 months
▫ Unsure
▫ Other (please specify) ________________________________________________

**C8.** For how long did you experience the new or worsening mood symptoms while having the arm implant?

☐ Less than 1 week
☐ 1 week - 1 month
☐ 1 - 3 months
☐ 3 - 6 months
☐ 6 months or more
☐ Unsure
☐ Other (please specify) ________________________________________________

**C9.** For how long did you have the arm implant after noticing the new or worsening mood symptoms?

☐ Less than 1 week
☐ 1 week - 1 month
☐ 1 - 3 months
☐ 3 - 6 months
☐ 6 months or more

- ☐ Unsure
- ☐ Other (please specify) ____________________________________________________

**C10.** Did the new/worsening mood symptoms you experienced after the arm implant was inserted go away when the implant was removed?

- ◻ Yes, completely
- ◻ Yes, partially
- ◻ No
- ◻ Not Applicable (please specify) ______________________________________________

  *[If yes]*

  After the arm implant was removed, how long did it take for the new/worsening mood symptoms to improve or go away?

  - ◻ Within a few days
  - ◻ Within 1 week
  - ◻ Within 1 month
  - ◻ Within 3 months
  - ◻ Within 6 months
  - ◻ Longer than 6 months
  - ◻ Unsure
  - ◻ Other (please specify) _____________________________________________

**C11.** Did you experience the above mood symptoms in response to more than one type/brand of arm implant?

- ◻ Yes
- ◻ No

---

*If Intrauterine device (IUD):*

**C1.** Which of the following symptoms started or became worse after the IUD was inserted? How severe were these symptoms? If you have experienced mood symptoms in response to more than one type/brand of IUD, respond below thinking of the type/brand that caused the most severe or most consistent mood symptoms.

| **Symptom** | **Not at all** | **Mild** | **Moderate** | **Severe** |
|---|---|---|---|---|
| 1. Anger/irritability | | | | |
| 2. Anxiety/tension | | | | |
| 3. Tearful/increased sensitivity to rejection | | | | |
| 4. Depressed mood/hopelessness | | | | |
| 5. Decreased interest in work activities | | | | |
| 6. Decreased interest in home activities | | | | |

| |
|---|
| 7. Decreased interest in social activities |
| 8. Difficulty concentrating |
| 9. Fatigue/lack of energy |
| 10. Overeating/food cravings |
| 11. Insomnia |
| 12. Hypersomnia (needing more sleep) |
| 13. Feeling overwhelmed or out of control |
| 14. Physical symptoms: breast tenderness, headaches, joint/muscle pain, bloating, weight gain |

**C2.** Did the symptoms you experienced interfere with:

| Symptom | Not at all | Mild | Moderate | Severe |
|---|---|---|---|---|
| A. Your work efficiency or productivity | | | | |
| B. Your relationships with coworkers | | | | |
| C. Your relationships with your family | | | | |
| D. Your social life activities | | | | |
| E. Your home responsibilities | | | | |

**C3.** If you know which type/brand of IUD led to these symptoms, please specify it here. If you have experienced mood symptoms in response to more than one type/brand of IUD, please provide the type/brand of the IUD that caused the most severe or most consistent mood symptoms. *If possible, please check your prescription history, packaging, or medication records before answering.*

☐ Yes

______________________________________________________________

☐ I don't know

*[If any response except I don't know]*

How confident are you in your answer about which IUD(s) led to these symptoms?

☐ I confirmed this using medical records, prescription history, or packaging
☐ Very confident — I clearly remember the exact IUD or brand
☐ Moderately confident — I think this is correct but am not completely sure
☐ Slightly confident — this is my best guess
☐ Not confident — I am unsure which IUD it was

**C4.** Please select the type of IUD that led to these symptoms.

☐ Hormonal
☐ Copper
☐ I don't know

*[If any response except I don't know]*
How confident are you in your answer about the type of IUD that led to these symptoms?
☐ I confirmed this using medical records, prescription history, or packaging
☐ Very confident — I clearly remember the exact IUD or brand
☐ Moderately confident — I think this is correct but am not completely sure
☐ Slightly confident — this is my best guess
☐ Not confident — I am unsure which IUD it was

**C5.** For how long did you have this IUD?
☐ Less than 1 week
☐ 1 week - 1 month
☐ 1 - 3 months
☐ 3 - 6 months
☐ 6 months or more
☐ Unsure

**C6.** How long after this IUD was inserted did you first notice new or worsening mood symptoms?
◻ Within 1 day
◻ Within 1 week
◻ Within 1 month
◻ Within 3 months
◻ Within 6 months
◻ Longer than 6 months
◻ Unsure
◻ Other (please specify) ____________________________________________

**C7.** For how long did you experience the new or worsening mood symptoms while having the IUD?
☐ Less than 1 week
☐ 1 week - 1 month
☐ 1 - 3 months
☐ 3 - 6 months
☐ 6 months or more
☐ Unsure
☐ Other (please specify) ____________________________________________

**C8.** For how long did you have the IUD after noticing the new or worsening mood symptoms?
☐ Less than 1 week

☐ 1 week - 1 month
☐ 1 - 3 months
☐ 3 - 6  months
☐ 6 months or more
☐ Unsure
☐ Other (please specify) ___________________________________________

**C9.** Did the new/worsening mood symptoms you experienced after the IUD was inserted get better when the IUD was removed?

□ Yes, completely
□ Yes, partially
□ No
□ Not Applicable (please specify) ______________________________________

*[If yes]*

After the IUD was removed, how long did it take for the new/worsening mood symptoms to improve or resolve? In other words, how long did it take for your mood to return to the same level as before the IUD was inserted?

□ Within 1 day
□ Within 1 week
□ Within 1 month
□ Within 3 months
□ Within 6 months
□ Longer than 6 months
□ Unsure
□ Other (please specify) _________________________________________

---

*If Injectable contraceptive (e.g. birth control shot, DepoProvera):*

**C1.** Which of the following symptoms started or became worse after starting the injectable contraceptive?  How severe were these symptoms when you were using the injectable contraceptive? If you have experienced mood symptoms in response to more than one type/brand of injectable contraceptive, respond below thinking of the type/brand of injectable contraceptive that caused the most severe or most consistent mood symptoms.

| **Symptom** | **Not at all** | **Mild** | **Moderate** | **Severe** |
|---|---|---|---|---|
| 1. Anger/irritability | | | | |
| 2. Anxiety/tension | | | | |
| 3. Tearful/increased sensitivity to rejection | | | | |
| 4. Depressed mood/hopelessness | | | | |
| 5. Decreased interest in work activities | | | | |

| |
|---|
| 6. Decreased interest in home activities |
| 7. Decreased interest in social activities |
| 8. Difficulty concentrating |
| 9. Fatigue/lack of energy |
| 10. Overeating/food cravings |
| 11. Insomnia |
| 12. Hypersomnia (needing more sleep) |
| 13. Feeling overwhelmed or out of control |
| 14. Physical symptoms: breast tenderness, headaches, joint/muscle pain, bloating, weight gain |

**C2.** Did the symptoms you experienced interfere with:

| Symptom | Not at all | Mild | Moderate | Severe |
|---|---|---|---|---|
| A. Your work efficiency or productivity | | | | |
| B. Your relationships with coworkers | | | | |
| C. Your relationships with your family | | | | |
| D. Your social life activities | | | | |
| E. Your home responsibilities | | | | |

**C3.** If you know which injectable contraceptive led to these symptoms, please provide it here. If you have experienced mood symptoms in response to more than one type/brand of injectable contraceptive, please provide the type/brand of the injectable that caused the most severe or most consistent mood symptoms. *If possible, please check your prescription history, packaging, or medication records before answering.*

☐ Yes

______________________________________________

☐ I don't know

*[If any response except I don't know]*
How confident are you in your answer about which injectable contraceptive(s) led to these symptoms?

☐ I confirmed this using medical records, prescription history, or packaging
☐ Very confident — I clearly remember the exact injectable contraceptive or brand
☐ Moderately confident — I think this is correct but am not completely sure
☐ Slightly confident — this is my best guess

☐ Not confident — I am unsure which injectable contraceptive it was

**C4.** How was the contraceptive injected?

- ☐ Intramuscular (into a muscle)
- ☐ Subcutaneous (under the skin)
- ☐ I don't know

**C5.** For how long did this injectable contraceptive provide contraception, from your first injection through the time covered by your last injection?

- ☐ Less than 3 months
- ☐ 3 months to less than 6 months
- ☐ 6 months to less than 12 months
- ☐ 12 months or longer
- ☐ Unsure
- ☐ Other (please specify) ______________________________________________

**C6.** How long after starting this injectable contraceptive did you first notice new or worsening mood symptoms?

- ☐ The day I got the first shot
- ☐ Within 1 week after getting the first shot
- ☐ Within 1 month after getting the first shot
- ☐ Within 3 months after getting the first shot
- ☐ Within 1 month of getting the second shot
- ☐ Within 3 months after getting the second shot
- ☐ After getting the third shot
- ☐ Unsure
- ☐ Other (please specify) ______________________________________________

**C7.** For how long did you experience the new or worsening mood symptoms while taking this injectable contraceptive?

- ☐ 1 week or less
- ☐ 1 week - 1 month
- ☐ 1 - 3 months
- ☐ 3 - 6 months
- ☐ 6 months or more
- ☐ Unsure
- ☐ Other (please specify) ______________________________________________

**C8.** How long did you continue taking this injectable contraceptive after noticing the new or worsening mood symptoms?

- ☐ Less than 1 week
- ☐ 1 week - 1 month

- ☐ 1 - 3 months
- ☐ 3 - 6 months
- ☐ 6 months or more
- ☐ Unsure
- ☐ Other (please specify) ___________________________________________________

**C9.** Did the new/worsening mood symptoms you experienced when starting this injectable contraceptive go away when you stopped taking this medication?

- ☐ Yes, completely
- ☐ Yes, partially
- ☐ No
- ☐ Not Applicable (please specify) _____________________________________________

  *[If yes]*

  After your last injectable contraceptive, how long did it take for the new/worsening mood symptoms to improve or go away?

  - ☐ Within 1 week after getting the last shot
  - ☐ Within 1 month after getting the last shot
  - ☐ Within 3 months after getting the last shot
  - ☐ Within 6 months after getting the last shot
  - ☐ Longer than 6 months after getting the last shot
  - ☐ Unsure
  - ☐ Other (please specify) ______________________________________________

**C10.** Did you experience the above mood symptoms in response to more than one type/brand of injectable contraceptive?

- ☐ Yes
- ☐ No

*If Fertility treatments (e.g. Clomid, Letrozole, Ovidrel, hCG, FSH, etc):*

**C1.** Which of the following symptoms started or became worse while taking hormonal medications for infertility? How severe were these symptoms? Only include symptoms that started and/or worsened within a few days of taking hormonal medications for infertility.

| **Symptom** | **Not at all** | **Mild** | **Moderate** | **Severe** |
|---|---|---|---|---|
| 1. Anger/irritability | | | | |
| 2. Anxiety/tension | | | | |
| 3. Tearful/increased sensitivity to rejection | | | | |
| 4. Depressed mood/hopelessness | | | | |
| 5. Decreased interest in work activities | | | | |

| |
|---|
| 6. Decreased interest in home activities |
| 7. Decreased interest in social activities |
| 8. Difficulty concentrating |
| 9. Fatigue/lack of energy |
| 10. Overeating/food cravings |
| 11. Insomnia |
| 12. Hypersomnia (needing more sleep) |
| 13. Feeling overwhelmed or out of control |
| 14. Physical symptoms: breast tenderness, headaches, joint/muscle pain, bloating, weight gain |

**C2.** Did the symptoms you experienced interfere with:

| Symptom | Not at all | Mild | Moderate | Severe |
|---|---|---|---|---|
| A. Your work efficiency or productivity | | | | |
| B. Your relationships with coworkers | | | | |
| C. Your relationships with your family | | | | |
| D. Your social life activities | | | | |
| E. Your home responsibilities | | | | |

**C3**. If you know which medication(s) led to these symptoms, please specify them here. Check all that apply. If possible, please check your prescription history, packaging, or medication records before answering.

☐ Clomiphene (Clomid)
☐ Letrozole (Femara)
☐ Choriogonadotropin alfa (Ovidrel)
☐ hCG
☐ FSH
☐ Other (please specify) ____________________________________
☐ I don't know

*[If any response except I don't know]*
How confident are you in your answer about which medication(s) led to these symptoms?
☐ I confirmed this using medical records, prescription history, or packaging
☐ Very confident — I clearly remember the exact medication or brand
☐ Moderately confident — I think this is correct but am not completely sure

☐ Slightly confident — this is my best guess
☐ Not confident — I am unsure which medication it was

**C4.** For how many cycles did you use one of the above medications?
☐ 1
☐ 2 - 3
☐ 4 - 6
☐ 7+

**C5.** In how many cycles did you experience new/worsening mood symptoms that started within a few days of starting one of the above medications and improved within a few days of stopping the medication?
☐ 1
☐ 2 - 3
☐ 4 - 6
☐ 7+

**C6.** How long after starting one of the above medications did you first notice new or worsening mood symptoms?
☐ 1 day
☐ 2 - 3 days
☐ 4 days–1 week
☐ 1 - 2 weeks
☐ 2 - 4 weeks
☐ 4+ weeks

**C7.** Within one cycle, how long on average did you experience the new or worsening mood symptoms while taking fertility medications?
☐ 1 day
☐ 2 - 3 days
☐ 4 days - 1 week
☐ 1 - 2 weeks
☐ 2 - 4 weeks
☐ 4+ weeks
☐ Other (please specify) ____________________________________________

**C8.** Did the new/worsening mood symptoms you experienced when starting fertility medication go away when you stopped taking the medication?
□ Yes, completely
□ Yes, partially
□ No
□ Not Applicable (please specify) ________________________________________
*[If yes]*

After you stopped taking the medication, how long did it take for the new/worsening mood symptoms to improve or go away?

- ☐ Within a few days
- ☐ Within 1 week
- ☐ Within 1 month
- ☐ Within 3 months
- ☐ Within 6 months
- ☐ Longer than 6 months
- ☐ Unsure
- ☐ Other (please specify) ________________________________________

### *Family History*

**4.** Have any of your first-degree relatives (biological mother or full biological siblings) experienced perinatal depression? Perinatal depression (sometimes referred to as postpartum depression) is new depression that started during pregnancy or within 12 months after delivery OR depression that became worse during pregnancy or within 12 months after delivery. Do not include depression that was present before pregnancy and continued at the same level during pregnancy. Do not include short-term "baby blues" that resolved within about 2 weeks after delivery.

☐ Yes
☐ No
☐ I don't know

*If "Yes":*

**C1.** Who in your family has experienced perinatal depression? Perinatal depression (sometimes referred to as postpartum depression) is new depression that started during pregnancy or within 12 months after delivery OR depression that became worse during pregnancy or within 12 months after delivery. Do not include depression that was present before pregnancy and continued at the same level during pregnancy. Do not include short-term "baby blues" that resolved within about 2 weeks after delivery. Check all that apply

☐ Biological mother
☐ Full biological sibling(s)

How many of your full biological siblings have experienced perinatal depression?

________________________________________________

---

*If Mother:*

During how many pregnancies did your mother experience perinatal depression?

☐ 1 ☐ 2 ☐ 3 ☐ 4 ☐ 5 ☐ 6 ☐ 7 ☐ 8 or more ☐ Unsure

Which of the following best describes your mother's depression for the pregnancy during which her perinatal depression was most severe?

☐ She was not actively depressed just before that pregnancy, and a new depressive episode began during pregnancy
☐ She was not actively depressed just before that pregnancy or during pregnancy, and a new depressive episode began after delivery
☐ She was already actively depressed just before that pregnancy, and her depression became worse during pregnancy
☐ She was already actively depressed before pregnancy, and her depression became worse after delivery
☐ None of the above - She was depressed before pregnancy, but the depression did not clearly worsen during pregnancy or after delivery

---

☐ Other (please specify): ___________________________
☐ Not sure

How do you know the information you reported about your mother's perinatal depression? (Check all that apply.)
☐ My mother told me directly about her experience
☐ Another family member told me
☐ I was present during the pregnancy or postpartum period and observed symptoms
☐ A healthcare provider told me
☐ I verified this using medical records
☐ I inferred or assumed based on what I remember
☐ Other (please specify): _______

To your knowledge, was your mother ever diagnosed by a healthcare professional with postpartum depression or perinatal depression for any pregnancy?
☐ Yes
☐ No
☐ Not sure

Overall, how confident are you in the information you provided about your mother's perinatal depression?
☐ Very confident
☐ Moderately confident
☐ Slightly confident
☐ Not confident

---

*If Full sibling:*
During how many pregnancies did your sibling experience perinatal depression?
☐ 1 ☐ 2 ☐ 3 ☐ 4 ☐ 5 ☐ 6 ☐ 7 ☐ 8 or more ☐ Unsure

Which of the following best describes your sibling's depression for the pregnancy during which her perinatal depression was most severe?
☐ She was not actively depressed just before that pregnancy, and a new depressive episode began during pregnancy
☐ She was not actively depressed just before that pregnancy or during pregnancy, and a new depressive episode began after delivery
☐ She was already actively depressed just before that pregnancy, and her depression became worse during pregnancy
☐ She was already actively depressed before pregnancy, and her depression became worse after delivery
☐ None of the above - She was depressed before pregnancy, but the depression did not clearly worsen during pregnancy or after delivery
☐ Other (please specify): ___________________________
☐ Not sure

How do you know the information you reported about your sibling’s perinatal depression? (Check all that apply.)
☐ My sibling told me directly about her experience
☐ Another family member told me
☐ I was present during the pregnancy or postpartum period and observed symptoms
☐ A healthcare provider told me
☐ I verified this using medical records
☐ I inferred or assumed based on what I remember
☐ Other (please specify): _______

To your knowledge, was your sibling ever diagnosed by a healthcare professional with postpartum depression or perinatal depression for any pregnancy?
☐ Yes
☐ No
☐ Not sure

Overall, how confident are you in the information you provided about your sibling’s perinatal depression?
☐ Very confident
☐ Moderately confident
☐ Slightly confident
☐ Not confident
