## Supplementary Material 2. Interval Depression Inventory for "The PMADS Project: A Longitudinal Multimodal Cohort Study to Understand Risk for Perinatal Mood and Anxiety Disorders"

### Interval Anxiety Inventory

**Instructions**

*The following questions ask about how you have been feeling* ***since your last study visit****. Please select the response that best reflects your experience during that time— not just how you feel today.*

**1.** Since your last study visit, has there been a period of time when you felt nervous, anxious, or on edge most of the day nearly every day?

☐ Yes ☐ No

*If Yes:*

**1a.** How long did that period last?

☐ Less than 1 week
☐ At least 1 week but less than 2 weeks
☐ At least 2 weeks but less than 4 weeks
☐ At least 4 weeks

**1b.** Approximately how many weeks ago did you start feeling nervous, anxious, or on edge?

☐ 1 ☐ 2 ☐ 3 ☐ 4 ☐ 5 ☐ 6 ☐ 7 ☐ 8 ☐ 9 ☐ 10 ☐ 11 ☐ 12 ☐ >12

**2.** Since your last study visit, has there been a period of time when you could not stop worrying for most of the day nearly every day?

☐ Yes ☐ No

*If Yes:*

**2a**. How long did that period last?
☐ Less than 1 week
☐ At least 1 week but less than 2 weeks
☐ At least 2 weeks but less than 4 weeks
☐ At least 4 weeks

**2b.** Approximately how many weeks ago did you start worrying for most of the day?
☐ 1 ☐ 2 ☐ 3 ☐ 4 ☐ 5 ☐ 6 ☐ 7 ☐ 8 ☐ 9 ☐ 10 ☐ 11 ☐ 12 ☐ >12

**3.** Have you ever been diagnosed with an anxiety disorder?

☐ Yes ☐ No

*If Yes:*

**3a**. Since your last study visit, has there been a period of time when your anxiety symptoms returned or worsened compared to your last visit?

☐ Yes ☐ No

*If Yes:*

**3b**. How long did that period last?

☐ Less than 1 week
☐ At least 1 week but less than 2 weeks
☐ At least 2 weeks but less than 4 weeks
☐ At least 4 weeks

**4.** Have you ever received treatment for anxiety symptoms (for example, therapy or medications)?

☐ Yes ☐ No

*If Yes:*

**4a.** Have you ever been prescribed medication(s) to treat anxiety symptoms?

☐ Yes ☐ No

**5.** Since your last study visit, have there been any changes in medications you are or were taking to treat anxiety symptoms? (Select all that apply.)

☐ Yes – I started taking a new medication for anxiety and am currently taking it
☐ Yes – I stopped taking a medication for anxiety that I was taking at my last study visit
☐ Yes – The total daily dose of a medication I am currently taking for anxiety was increased
☐ Yes – The total daily dose of a medication I am currently taking for anxiety was decreased
☐ Yes – I started taking a new medication for anxiety since my last study visit but am no longer taking it
☐ Yes – A provider prescribed a new medication for anxiety, but I did not take it
☐ Yes – Other medication change not listed above
☐ No – There have been no changes in anxiety medications
☐ No – I have not taken any medications for anxiety since my last study visit

*If you selected any "Yes" or "Other," please specify:*

**6.** Since your last study visit, have there been any changes in non-medication treatments you are or were receiving for anxiety? (Select all that apply.)

☐ Yes – I started seeing a therapist and am currently seeing a therapist
☐ Yes – I started seeing a therapist but am not currently seeing one
☐ Yes – I am seeing a therapist more frequently than at my last study visit
☐ Yes – I started seeing an additional therapist
☐ Yes – I started seeing an additional therapist but am no longer seeing them
☐ Yes – I temporarily increased therapy frequency but am no longer doing so
☐ Yes – I completed or am currently in an intensive outpatient program
☐ Yes – I completed a partial hospitalization program
☐ Yes – I was admitted to an inpatient psychiatric hospital
☐ Yes – I received another treatment for anxiety not listed here (for example, transcranial magnetic stimulation, infusion-based treatments)
☐ No – There have been no changes in non-pharmacologic anxiety treatments
☐ No – I have not received any non-pharmacologic treatment for anxiety since my last study visit

**If you selected any "Yes" or "Other," please specify:**

______________________________________________

______________________________________________
