## Supplementary Material 3. Interval Anxiety Inventory for "The PMADS Project: A Longitudinal Multimodal Cohort Study to Understand Risk for Perinatal Mood and Anxiety Disorders"

### Interval Depression Inventory

**Instructions**

*The following questions ask about how you have been feeling **since your last study visit**. Please select the response that best reflects how you've felt during that time— not just today.*

**1.** Since your last study visit, has there been a period of time when you were feeling down or depressed most of the day nearly every day?

☐ Yes ☐ No

*If Yes:*

**1a.** How long did that period last?

☐ Less than 1 week
☐ At least 1 week but less than 2 weeks
☐ At least 2 weeks but less than 4 weeks
☐ At least 4 weeks

**1b.** Approximately how many weeks ago did you start feeling down or depressed?

☐ 1 ☐ 2 ☐ 3 ☐ 4 ☐ 5 ☐ 6 ☐ 7 ☐ 8 ☐ 9 ☐ 10 ☐ 11 ☐ 12 ☐ >12

**2.** *If you answered **No** to Question 1, please respond to question 2. Otherwise, please skip to Question 3*

Since your last study visit, has there been a period of time when you felt empty or hopeless most of the day nearly every day?

☐ Yes ☐ No

*If Yes:*

**2a**. How long did that period last?
☐ Less than 1 week
☐ At least 1 week but less than 2 weeks
☐ At least 2 weeks but less than 4 weeks
☐ At least 4 weeks

**2b.** Approximately how many weeks ago did you start feeling empty or hopeless?
☐ 1 ☐ 2 ☐ 3 ☐ 4 ☐ 5 ☐ 6 ☐ 7 ☐ 8 ☐ 9 ☐ 10 ☐ 11 ☐ 12 ☐ >12

Since your last study visit, has anyone said that you look sad, down, or depressed*?*

☐ Yes ☐ No

*If Yes:*

**2a**. How long did that period last?
☐ Less than 1 week
☐ At least 1 week but less than 2 weeks
☐ At least 2 weeks but less than 4 weeks
☐ At least 4 weeks

**2b.** Approximately how many weeks ago did you start feeling down or depressed?
☐ 1 ☐ 2 ☐ 3 ☐ 4 ☐ 5 ☐ 6 ☐ 7 ☐ 8 ☐ 9 ☐ 10 ☐ 11 ☐ 12 ☐ >12

**3.** Since your last study visit, has there been a period of time when you lost interest or pleasure in almost all activities you usually enjoyed?

☐ Yes ☐ No

*If Yes:*

**3a**. How long did that period last?

☐ Less than 1 week
☐ At least 1 week but less than 2 weeks
☐ At least 2 weeks but less than 4 weeks
☐ At least 4 weeks

**2b.** Approximately how many weeks ago did you start feeling down or depressed?
☐ 1 ☐ 2 ☐ 3 ☐ 4 ☐ 5 ☐ 6 ☐ 7 ☐ 8 ☐ 9 ☐ 10 ☐ 11 ☐ 12 ☐ >12

**4.** Have you ever been diagnosed with depression?

☐ Yes ☐ No

*If Yes:*

**4a**. Since your last study visit, has there been a period of time where you noticed that your depressive symptoms returned or worsened (compared to the level they were at during your last visit)?

☐ Yes ☐ No

*If Yes:*

**4b**. How long did that period last?

☐ Less than 1 week
☐ At least 1 week but less than 2 weeks
☐ At least 2 weeks but less than 4 weeks
☐ At least 4 weeks

**4c.** Approximately how many weeks ago did you start feeling down or depressed?
☐ 1 ☐ 2 ☐ 3 ☐ 4 ☐ 5 ☐ 6 ☐ 7 ☐ 8 ☐ 9 ☐ 10 ☐ 11 ☐ 12 ☐ >12

**5.** Have you ever received treatment for symptoms of depression (for example, therapy or medications)?

☐ Yes ☐ No

*If Yes:*

**5a.** Have you ever been prescribed medication(s) to treat symptoms of depression?

☐ Yes ☐ No

**6.** *If you answered **Yes** to Question 5a*

Since your last study visit, have there been any changes in medications you are or were taking to treat symptoms of depression? (Select all that apply.)

- ☐ Yes – I have started taking a new medication for depression since my last study visit
- ☐ Yes – I stopped taking a medication for depression that I was taking at my last study visit
- ☐ Yes – The total daily dose of the medication I am currently taking for depression was increased
- ☐ Yes – The total daily dose of the medication I am currently taking for depression was decreased
- ☐ Yes – I started taking a new medication for depression since my last study visit but am no longer taking it
- ☐ Yes – A provider prescribed a new medication for depression but I have not taken it
- ☐ Yes – Other change in medication not listed here
- ☐ No – There have been no changes in medications I am taking for depression
- ☐ No – I have not taken any medications for depression since my last study visit
- ☐ Other

*If you selected any "Yes" or "Other," please specify:*

Since your last study visit, have there been any changes in treatments other than medication you are or were receiving to treat symptoms of depression? (Select all that apply.)

- □ Yes – I started seeing a therapist and am currently seeing a therapist
- □ Yes – I started seeing a therapist but am not currently seeing a therapist
- □ Yes – I am seeing a therapist(s) more frequently than I was at the time of my last study visit
- □ Yes – I have started seeing an additional therapist
- □ Yes – I started seeing an additional therapist but I am no longer seeing the additional therapist
- □ Yes – There was a period of time where I was seeing a therapist more frequently than I was at the time of my last study visit but I am no longer seeing them at an increased frequency
- □ Yes – I completed or am in an intensive outpatient program
- □ Yes – I completed a partial hospitalization program
- □ Yes – I was admitted to an inpatient psychiatric hospital
- □ Yes – I received a treatment for depression not listed here (for example, transcranial magnetic stimulation, electroconvulsive therapy, ketamine, infusion-based treatments etc.)
- □ No – there have been no changes in non-pharmacologic treatments I am receiving for depression
- □ No – I have not received any non-pharmacologic treatments for depression since my last study visit
- □ Other

*If you selected any "Yes" or "Other," please specify:*

________________________________________

________________________________________
